# Computational investigation of circuit mechanisms underlying short-latency responses to cortical stimulation

**DOI:** 10.64898/2026.09.11.750854

**Authors:** Karthik Kumaravelu, Gene J. Yu, Aman S. Aberra, Marc A. Sommer, Angel V. Peterchev, Warren M. Grill

## Abstract

Transcranial magnetic stimulation (TMS) over the primary motor cortex (M1) elicits a series of high frequency volleys termed D- and I-waves measured epidurally in the corticospinal tract of awake humans. Further, intracortical microstimulation (ICMS) in M1 of non-human primates evokes D- and I-wave responses similar to those observed in TMS. The cortical circuits and mechanisms involved in the generation of D- and I-waves by stimulation of M1 remain unclear. Here, we implemented computational models of cortical columns with laminarly-organized biophysically-based neurons, following existing models published in the literature: (1) M1 - single compartment (SC), (2) M1 - multi-compartment (MC), (3) primary auditory cortex (A1) - MC, and (4) primary somatosensory cortex (S1) - MC. The network connectivity of each model represented wiring found in the respective cortical regions. The direct effects of stimulation-induced electric fields were modeled as activation of different proportions of pyramidal neurons (PNs) across layers, and dose–response curves were constructed for layer 5 (L5) PNs. Both the M1 and A1 models reproduced D- and I-waves, with the magnitude of I-waves increasing with higher recruitment of layer 2/3 and layer 5 PNs. The S1-MC model evoked rhythmic firing activity but with timings mismatched to experimental I-waves. The models replicated the experimentally observed effects of pharmacological agents on I-waves, and virtual lesions of specific neural populations across models revealed plausible microcircuit explanations for the first and later I-waves. This comprehensive comparison of models across multiple cortical regions identified consistent mechanisms underlying the cortical response to TMS and contributes to the refinement of computational strategies for optimizing stimulation paradigms.

**Significance Statement:** Cortical stimulation modalities such as transcranial magnetic stimulation and intracortical microstimulation applied to the primary motor cortex generate highly stereotyped, short-latency responses termed D- and I-waves, measured epidurally in the corticospinal tract of humans and non-human primates. Direct recording of the TMS- and ICMS-evoked neural response in the cortex is extremely challenging due to stimulation-induced artifacts. Furthermore, the short-latency responses evoked by stimulation of non-motor cortical regions remain unknown. Using sophisticated computational models representing different cortical areas, we simulated the immediate responses to cortical stimulation modalities including TMS and ICMS to characterize the circuit mechanisms underlying D- and I-waves. Gaining a better understanding of the cortical circuit mechanisms enables the interpretation and optimization of stimulation paradigms.

## INTRODUCTION

Transcranial magnetic stimulation (TMS) applied to the primary motor cortex (M1) can elicit motor evoked potentials (MEPs) in target muscles. This technique is used to quantify cortical excitability and individualize TMS treatments (Paulus et al. 2013). TMS MEPs are driven by the corticospinal tract (CST) response, which comprises a series of high frequency (∼600 Hz), short-latency (1–6 ms) volleys termed D- (for direct) and I- (for indirect) waves measured in awake humans (Di Lazzaro et al. 1998, Di Lazzaro et al. 2012, Di Lazzaro & Rothwell 2014, Di Lazzaro & Ziemann 2013, Di Lazzaro et al. 2008, Ziemann & Rothwell 2000) and in anesthetized non-human primates (NHPs) (Edgley et al. 1997, Glover et al. 2026). Similarly, with recent methodological improvements to mitigate artifacts, EEG during TMS over M1 in human subjects showed several short-latency peaks resembling I-wave activity (Beck et al. 2024, Nuyts et al. 2025). As well, intracortical microstimulation (ICMS) in M1 of NHPs evokes D- and I-waves similar to those evoked by TMS (Lemon 2008, Maier et al. 2013, Shimazu et al. 2004). The D-wave is produced by *direct* activation by the stimulation-induced electric field of CST axons originating in layer 5 (L5) of the motor cortex, while the longer latency I-waves are thought to be produced by repetitive *indirect* activation of CST neurons through trans-synaptic excitation (Di Lazzaro & Ziemann 2013, Di Lazzaro et al. 2008). However, the microcircuit mechanisms underlying the generation of I-waves remain unclear.

I-waves exhibit several hallmark characteristics (Di Lazzaro et al. 1998, Di Lazzaro et al. 2012, Di Lazzaro & Ziemann 2013, Di Lazzaro et al. 2008, Esser et al. 2005, Rusu et al. 2014). First, I-waves increase in both amplitude and number as the intensity of TMS or ICMS is increased. Second, the magnitude of late I-waves diminishes after the application of GABA-A (GABA_A_R) receptor positive allosteric modulators (Burke et al. 1993, Di Lazzaro et al. 2000), implicating involvement of inhibitory GABAergic interneurons in generating late I-waves. Finally, the magnitude of I-waves is increased during increased cortical excitability such as during voluntary muscle contraction (Di Lazzaro 1998). However, despite decades of research, the neural origin and the cortical circuits responsible for generating I-waves have been elusive due to large stimulation artifacts during cortical recordings and the inability to selectively deactivate or lesion specific pathways (Di Lazzaro et al. 2012). Several theories exist regarding the origin of the I-wave response to TMS over M1, including both non-synaptic (cellular) and trans-synaptic (circuit) mechanisms (Di Lazzaro et al. 2012, Massimini et al. 2026, Siebner et al. 2022, Ziemann & Rothwell 2000). Although computational models have been used to simulate I-wave responses to M1 stimulation (Esser et al. 2005, Haggie et al. 2024, Rusu et al. 2014, Schaworonkow & Triesch 2018, Yen et al. 2012, Yu et al. 2024), there is no consensus mechanism, and there has been no investigation of short-latency responses evoked by stimulation applied over other cortical areas (for example, the somatosensory cortex).

This study aimed to explore neural sources that can contribute to the short-latency responses to M1 stimulation as well as non-motor cortical areas using computational modeling. The primary objective of this study was not to model the direct neuronal activation produced by TMS- or ICMS-induced E-fields (Aberra et al. 2020, Kumaravelu et al. 2022, Worbs et al. 2026), but rather to investigate, using detailed cortical network models, the subsequent network activity arising from synaptic propagation following that initial activation. We implemented established cortical column network models representing different cortical regions of the rodent brain (Borges et al. 2022, Dura-Bernal et al. 2023, Esser et al. 2005, Traub et al. 2005). We used several existing models from the literature to evaluate whether the experimentally observed short-latency responses arise as emergent properties of the models, rather than being reproduced through model fitting. We modeled the direct effects of the stimulation-induced electric fields as activation of pyramidal neurons (PNs), and L5 axonal spiking activity was used as the primary output measure. In the M1 models, this output serves as a proxy for the response of CST axons to stimulation. We quantified dose–response relationships across models and characterized the effects of pharmacological agents and synaptic pathway lesioning on the short-latency responses. The results reveal neural circuit mechanisms that can contribute to the short-latency responses to stimulation modalities including TMS and ICMS in different cortical areas.

## METHODS

We used four published models of interconnected networks of cortical neurons representing different cortical regions to study the short-latency neural responses to stimulation (Table 1): (1) network model of the motor cortex with single-compartment neurons (M1-SC), (2) network model of the motor cortex with multi-compartment neurons (M1-MC), (3) network model of the auditory cortex with multi-compartment neurons (A1-MC) and (4) network model of the somatosensory cortex with multi-compartment neurons (S1-MC). The data used to validate each model in their original publications are described in Table 2.

**Table 1:** Characteristics of the four models used in this study. M1-SC – model of motor cortex with single compartment neurons, M1-MC – model of motor cortex with multi-compartment neurons, A1-MC – model of auditory cortex with multi-compartment neurons, S1-MC – model of somatosensory cortex with multi-compartment neurons. PN – pyramidal neuron, IN – interneuron, PTN – pyramidal tract neuron, SS – spiny stellate neuron, CT – corticothalamic neuron.

| Model | Cortical region | Morphology | Neurons/layer | Synapse type |
| --- | --- | --- | --- | --- |
| M1-SC (Esser et al. 2005) | Rat M1 | Single compartment | L2/3: 6400 PNs, 3200 INs<br>L5: 6400 PTNs, 3200 INs<br>L6: 6400 PNs, 3200 INs | Chemical: AMPAR, NMDAR, GABA<sub>A</sub>R, GABA<sub>B</sub>R |
| M1-MC (Dura-Bernal et al. 2023) | Mouse M1 | Multi-compartment | L2/3: 62 PNs, 20 INs<br>L4: 35 PNs<br>L5A: 22 PNs and 9 INs<br>L5B: 41 PNs, 41 PTNs and 21 INs.<br>L6: 52 PNs, 52 CTs and 10 INs. | Chemical: AMPAR, NMDAR, GABA<sub>A</sub>R |
| A1-MC (Traub et al. 2005) | Rodent A1 | Multi-compartment | L2/3: 105 PNs, 27 INs<br>L4: 24 SS<br>L5: 100 PNs<br>L6: 50 PNs, 30 Ins | Chemical: AMPAR, NMDAR, GABA<sub>A</sub>R<br>Electrical: Gap junctions |
| S1-MC (Borges et al. 2022) | Rat S1 | Multi-compartment | L2/3: 5877 PNs, 1819 INs<br>L4: 2674 PNs/SS, 584 INs<br>L5: 5050 PNs, 2628 INs<br>L6: 4812 PNs and 7902 Ins | Chemical: AMPAR, NMDAR, GABA<sub>A</sub>R, GABA<sub>B</sub>R |

**Table 2:** Details of model validation used in the original publications. M1-SC – model of motor cortex with single compartment neurons, M1-MC – model of motor cortex with multi-compartment neurons, A1-MC – model of auditory cortex with multi-compartment neurons, S1-MC – model of somatosensory cortex with multi-compartment neurons.

| Model | How was the model validated in the original publication |
| --- | --- |
| M1-SC (Esser et al. 2005) | 1. Firing rates of L5, L6 and thalamus neurons matched against experimental data |
| M1-MC (Dura-Bernal et al. 2023) | 1. Firing rates of L2/3 and L5 pyramidal neurons during quiet wakefulness and movement states matched against experimental data<br>2. L5 LFP spectral power during quiet wakefulness and movement states matched against experimental data<br>3. Firing rate of L5B pyramidal neurons to different levels of motor thalamus and noradrenaline inputs matched against experimental data |
| A1-MC (Traub et al. 2005) | 1. Presence of very fast oscillations superimposed on epileptiform field potentials and firing patterns of L4 spiny stellate cells during seizure-like events matched against experimental data |
| S1-MC (Borges et al. 2022) | 1. VPM thalamus synaptic responses in L4 and L5 pyramidal neurons matched against experimental recordings |

### Network models of motor cortex

Since I-waves are most characterized experimentally during M1 stimulation (Di Lazzaro et al. 2012, Lemon 2008), we implemented and simulated two network models based on rodent M1 to quantify the I-wave responses (Dura-Bernal et al. 2023, Esser et al. 2005). The Esser et al., model (M1-SC) comprised ∼33,000 single-compartment integrate-and-fire cortical neurons arranged in a 3-layered cortical slab with dimensions 5.5 mm × 5.5 mm × 2 mm (Fig. 1A). The network contained layer 2/3 (L2/3), layer 5 (L5), and layer 6 (L6) and each layer included 6400 PNs and 3200 interneurons (INs) (Fig. 1A). The model included both excitatory (AMPAR and NMDAR) and inhibitory synapses (containing GABA_A_R, GABA_B_R) with synaptic properties constrained based on experimental data (Fig. 2B). The biophysical properties of the cells and the network properties, including synaptic connection strengths, type, density and delay are described in detail in the original publication (Esser et al. 2005), and the firing behavior of model neuron types in each layer matched membrane potential traces and firing rates of cells recorded in vivo in M1.

**Figure 1:**
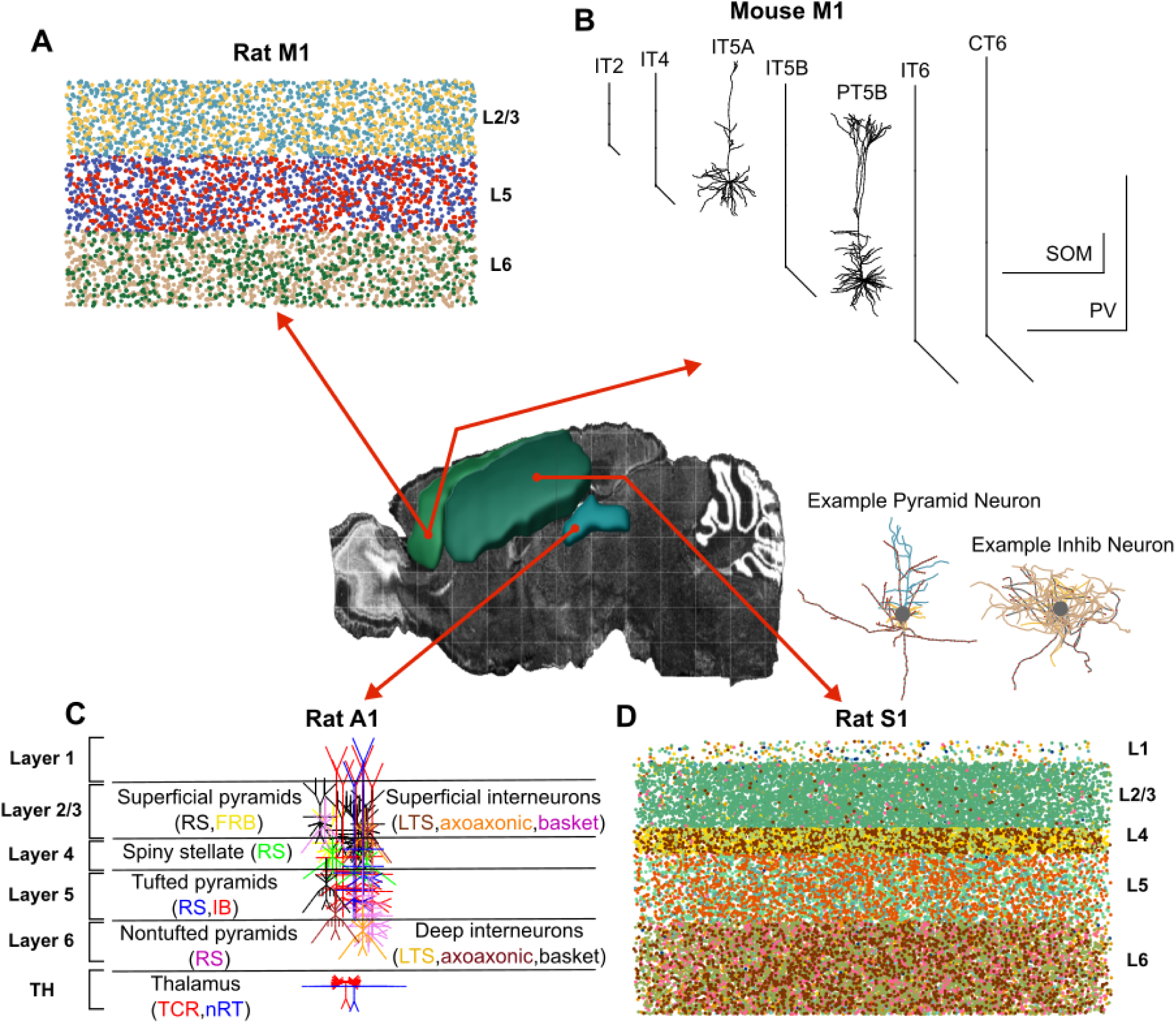
Different cortical regions, delineated on a sagittal slice of the rat brain, that were modeled to study the mechanisms of I-waves: (1) primary motor cortex (M1), (2) primary somatosensory cortex (S1) and (3) primary auditory cortex (A1). Biophysically-based computational models of different cortical regions. (A) M1-SC – model of motor cortex with single compartment neurons (Esser et al., 2005), (B) M1-MC – model of motor cortex with multi-compartment neurons (Dura-Bernal et al., 2023), (C) A1-MC – model of auditory cortex with multi-compartment neurons (Traub et al., 2005) and (D) S1-MC – model of somatosensory cortex with multi-compartment neurons (Borges et al., 2022). (A) The M1-SC model included 3200 E2/3(0) neurons, 3200 E2/3(180) neurons, 1600 I2/3(0) neurons, 1600 I2/3(180) neurons, 3200 E5(0) neurons, 3200 E5(180) neurons, 1600 I5(0) neurons, 1600 I5(180) neurons, 3200 E6(0) neurons, 3200 E6(180) neurons, 1600 I6(0) neurons, and 1600 I6(180) neurons arranged in a columnar format. ‘E’ and ‘I’ indicate excitatory and inhibitory neurons. Each neuron in the M1-SC model consisted of a single compartment (point neurons). The model consisted of neurons assigned to two opposing movement directions: 0 and 180 degrees, following the original Esser et al. model. (B) Morphology of cortical neurons across different layers in the M1-MC model. ‘IT’, ‘PT’ and ‘CT’ represent intratelencephalic, pyramidal tract and cortico-thalamic excitatory neurons. ‘SOM’ and ‘PV’ indicate somatostatin and parvalbumin inhibitory neurons. The M1-MC model included 62 IT2 neurons, 10 SOM2/PV2 neurons, 35 IT4 neurons, 22 IT5A neurons, 9 SOM5A/PV5A neurons, 41 IT5B neurons, 41 PT5B neurons, 21 SOM5B/PV5B neurons, 52 IT6 neurons, 52 CT6 neurons, and 10 SOM6/PV6 neurons arranged in a column. (C) Schematic of the cortical column showing the different populations of neurons across layers 1–6 (L1–L6) in the A1-MC model. Each color denotes a specific cell type characterized based on firing behavior. The cortical column model included 105 E2/3 pyramidal neurons, 27 I2/3 inhibitory interneurons, 24 E4 spiny stellate neurons, 100 E5 pyramidal neurons, 50 E6 pyramidal neurons, and 30 I6 inhibitory interneurons. The thalamus model included 10 thalamocortical relay neurons (TCR) and 10 reticular nucleus neurons (nRT). (D) Representative example of the morphology of a pyramidal neuron and an interneuron from the S1-MC model. The S1-MC model included 5877 E2/3 pyramidal neurons, 1819 I2/3 interneurons, 2674 E4 pyramidal and spiny stellate neurons, 584 I4 interneurons, 5050 E5 pyramidal neurons, 2628 I5 interneurons, 4812 E6 pyramidal neurons, and 7902 I6 interneurons arranged in a column.

**Figure 2:**
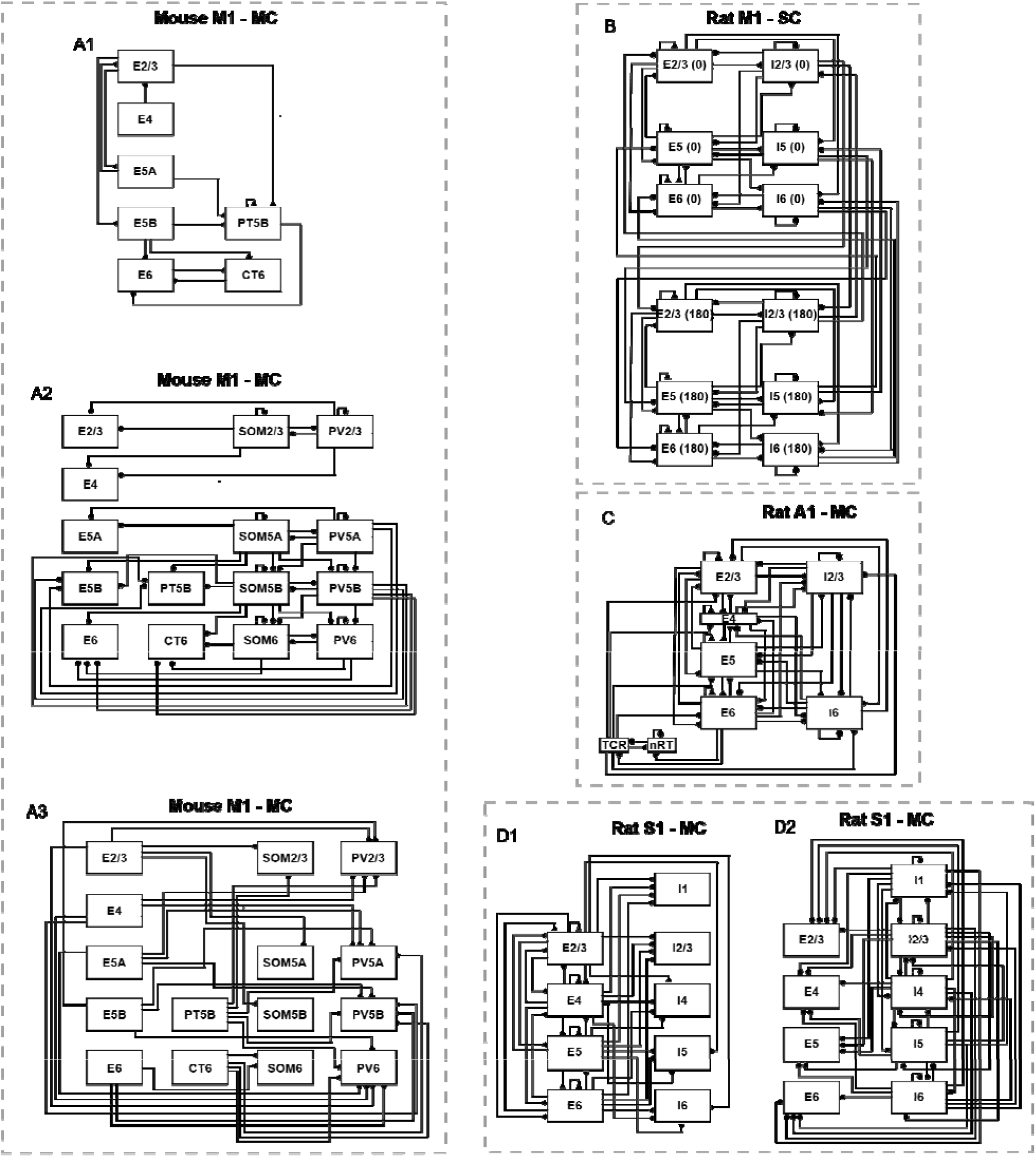
Synaptic connectivity across the four models. Synaptic connectivity of the M1-MC model showing (A1) excitatory inputs received by excitatory neurons, (A2) inhibitory inputs received by excitatory and inhibitory neurons, and (A3) excitatory inputs received by inhibitory neurons. Synaptic connectivity of (B) M1-SC model, (C) A1-MC model and (D) S1-MC model. Synaptic connectivity of the S1-MC model showing (D1) excitatory inputs received by excitatory and inhibitory neurons and (D2) inhibitory inputs received by excitatory and inhibitory neurons. The wiring diagrams show both excitatory (glutamate – AMPAR and NMDAR) and inhibitory (GABA_A_R and GABA_B_R) connections. For definitions, see caption of Figure 1.

Next, we implemented a second network model (more biophysically detailed than the M1-SC model) based on the mouse M1 to quantify the I-wave response (Dura-Bernal et al. 2022). The original Dura-Bernal et al., model (M1-MC) comprised 10,000 multi-compartment cortical neurons with ion channel dynamics based on the Hodgkin-Huxley model, i.e., Hodgkin-Huxley dynamics. However, due to the exceedingly high computational cost of the original model, we used a reduced model with 355 multi-compartment cortical neurons arranged in a column of six layers with dimensions of 400 µm × 400 µm × 1350 µm. The PNs from L5A and the pyramidal tract neurons (PTNs) from L5B were represented using detailed morphologies and ion channel properties, whereas simpler (2–6 compartment) models were used for other PNs and INs (Fig. 1B). Layer 1 (L1) contained tufted dendrites arising from the L5 PNs. L2/3 included 62 PNs and 10 somatostatin (SOM) expressing low-threshold spiking (LTS) and parvalbumin (PV) expressing fast spiking (FS) INs. L4 consisted of 35 PNs. L5A included 22 PNs and 9 INs, while L5B comprised 41 PNs, 41 PTNs and 21 INs. Finally, L6 included 52 PNs, 52 cortico-thalamic neurons (CTs) and 10 INs. Network interactions in the model were mediated through excitatory (AMPAR and NMDAR) and inhibitory (GABA_A_R) synapses (Fig. 2A1, A2, A3). The synaptic weights between connections in the reduced model were adjusted so that the reduced model replicated the emergent properties (such as firing rate, pattern, etc.) seen in the original model (Extended Data Fig. 2-1). Further details on the biophysical properties of the cells and the network properties can be found in the original publication (Dura-Bernal et al. 2023), and were primarily based on *in vivo* data from mouse M1.

**Figure 3:**
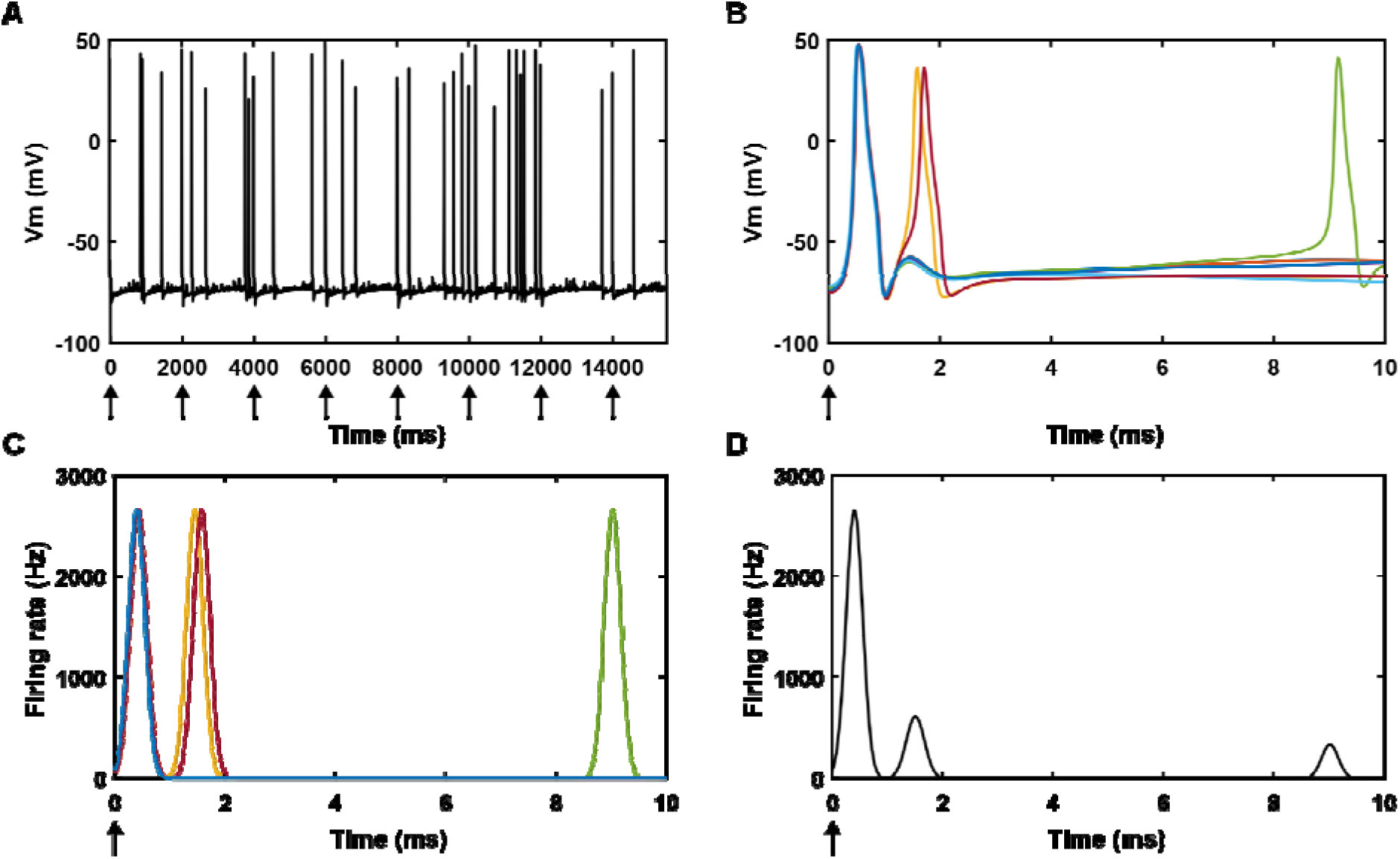
Response of an L5 pyramidal neuron (PN) to stimulation. Averaged spiking activity across L5 PNs was used as the model-based proxy for responses measured epidurally from the cortico-spinal tract of humans. (A) Transmembrane potential recorded from the axon terminal of an L5 PN. The stimulating pulse was applied at a frequency of 0.5 Hz (indicated by arrows). (B) Pulse-triggered responses shown across all eight stimulation pulses. Each color trace denotes the response to a different stimulation pulse. (C) Spike times were derived from the transmembrane potentials using threshold crossing and convolved with a Gaussian function to obtain smooth population firing rates across all stimulation pulses. Each color trace denotes the response to a different stimulation pulse. (D) Pulse-triggered response averaged across all stimulation pulses.

### Network model of auditory cortex

There are no experimental studies characterizing the I-wave response in sensory cortices such as the somatosensory cortex, auditory cortex, visual cortex etc. Additionally, the sensory cortices lack PNs that project directly to the spine like the PTNs of M1. For the sensory cortices, the L5 PNs were used to represent excitatory neurons that project to subcortical structures. To determine whether sensory cortices generate I-wave like responses, we implemented a computational model of the thalamo-cortical (TC) network using biophysically-based multi-compartment cortical neurons with Hodgkin-Huxley dynamics (Traub et al. 2005). The TC network model (A1-MC) included 356 neurons, each with 50–137 compartments. The cortex and thalamus were composed of twelve and two different populations of neurons, respectively, characterized by their firing properties. The cortical neurons were arranged in a columnar fashion consisting of six layers with a width of ∼400 µm and a depth of ∼2000 µm (Fig. 1C). L1 mostly contained the tufted dendrites arising from the L5 PNs. L2/3 included 100 regular spiking (RS) PNs, 5 fast rhythmic bursting (FRB) PNs, 18 FS INs and 9 LTS INs. L4 included 24 RS spiny stellate cells. L5 included 80 intrinsic bursting tufted PNs and 20 RS PNs. L6 included 50 RS non-tufted PNs, 20 FS INs and 10 LTS INs. The thalamic population consisted of 10 thalamocortical relay (TCR) cells and 10 reticular nuclei (nRT) cells. Network interactions in the model were mediated through chemical synapses (AMPAR, NMDAR and GABA_A_R) (Fig. 2C) and electrical gap junctions. The biophysical properties of the cells and the network properties, including synaptic connection strengths, type, density and delay are described in detail in the original publication (Traub et al. 2005), and were primarily based on *in vivo* and *in vitro* data from auditory cortex of rats.

The response to stimulation was quantified in the presence of the intrinsic gamma oscillatory activity occurring in the superficial layer (L2/3) in the original model. In addition to the intrinsic gamma oscillatory activity, the original model included ectopic spikes generated by injecting suprathreshold current pulses with inter-pulse intervals drawn from a Poisson distribution. The mean interval of the Poisson distribution was 10 s for superficial PNs and 1 s for all other excitatory neurons in the model.

The monosynaptic L2/3 to L5 excitatory connections are strong in M1 (Anderson et al. 2010), and it is hypothesized that TMS at motor threshold activates L2/3 PNs that, in turn, monosynaptically activate L5 PNs, generating the I_1_-wave (Di Lazzaro et al. 2012). However, in the Traub model, the L2/3 to L5 excitatory synapses were weak (Traub et al. 2005), and activation of L2/3 PNs in the original model did not produce any noticeable changes in the L5 PN activity. Therefore, we strengthened the connection between L2/3 to L5 PNs by making the connection density and strength the same as the recurrent connections from L5 PNs to L5 PNs (Extended Data Fig. 2-2).

### Network model of somatosensory cortex

We implemented a network model of the somatosensory cortex (S1-MC) based on rodent S1 developed as part of the Blue Brain project (Borges et al. 2022, Markram et al. 2015). The S1-MC model comprised 31,346 cells with detailed morphologies and Hodgkin-Huxley dynamics (55 different cortical neuron types characterized based on firing behavior and morphology) arranged in a columnar fashion consisting of six layers with a width of 420 µm and a depth of 2082 µm (Fig. 1D). L1 contained the tufted dendrites arising from the L5 PNs. L2/3 included 5877 PNs and 1819 INs. L4 comprised 2674 PNs/spiny stellate cells and 584 INs. L5 included 5050 PNs and 2628 INs, while L6 comprised 4812 PNs and 7902 INs. Network interactions in the model were mediated through chemical synapses (AMPAR, NMDAR, GABA_A_R and GABA_B_R) (Fig. 2D1, D2). The biophysical properties of the cells and the network properties, including synaptic connection strengths, type, densities, and delays are described in detail in the original publication (Borges et al. 2022, Markram et al. 2015), and were primarily based on *in vivo* and *in vitro* data from rat S1.

### Implementation of cortical stimulation

Prior efforts in modeling the network effects of TMS coupled the effects of TMS-induced electric field through either the activation of synapses or the direct activation of neurons via intracellular current injection (Table 3). We modeled the direct effects of stimulation-induced electric fields, such as those from TMS and ICMS, as activation of PNs in the superficial and deep layers (Table 4). To generate the response to repeated trials within single simulation runs, stimulation was delivered at a low frequency of 0.5 Hz (a total of 10 pulses) to avoid carry-over effects from previous pulses, similar to experimental single-pulse TMS and ICMS paradigms.

**Table 3:** Summary of how the TMS field was represented in other published network modeling studies. PN – pyramidal neuron, IN – interneuron, CST – corticospinal tract. MEP – motor evoked potential.

| Network Model | How was the TMS field represented | Additional information |
| --- | --- | --- |
| Esser et al. 2005 | Synaptic activation of randomly chosen PNs and INs across cortical layers |  |
| Yen et al. 2012 | Injecting a short pulse of direct current to 20% of randomly | Uses the Esser model but incorporates synaptic plasticity |
|  | chosen pre-synaptic terminals |  |
| Rusu et al. 2014 | Current injection into L2/3 soma and L5 axon initial segment |  |
| Schaworonkow & Triesch 2018 | Current injection into L2/3 soma and L5 axon initial segment | Uses the Rusu model but incorporates oscillatory activity in L2/3 PNs |
| Moezzi et al. 2018 | Current injection into L2/3 soma and L5 axon initial segment | Uses the Rusu model but incorporates the CST to model MEP |
| Haggie et al. 2024 | Direct activation of neurons with strength of activation linearly decreasing with depth |  |

**Table 4:** Summary of how the TMS field was represented across the four models in this study. M1-SC – model of motor cortex with single compartment neurons, M1-MC – model of motor cortex with multi-compartment neurons, A1-MC – model of auditory cortex with multi-compartment neurons, S1-MC – model of somatosensory cortex with multi-compartment neurons. PN – pyramidal neuron, IN – interneuron, PTN – pyramidal tract neuron.

| Model | How was the TMS field modeled | How was the I-wave response quantified |
| --- | --- | --- |
| M1-SC (Esser et al. 2005) | 1. Activation of L2/3 or L5 PNs<br>2. Simultaneous activation of L2/3 and L5 PNs | Spiking activity averaged across 6400 L5 PNs |
| M1-MC (Dura-Bernal et al. 2023) | 1. Activation of L2/3 PNs or (L5b PNs + L5b PTNs)<br>2. Simultaneous activation of L2/3 PNs and L5b PNs + L5b PTNs | Spiking activity averaged across 41 L5b PTNs |
| A1-MC (Traub et al. 2005) | 1. Activation of L2/3 or L5 PNs<br>2. Simultaneous activation of L2/3 and L5 PNs | Spiking activity averaged across 100 L5 PNs |
| S1-MC (Borges et al. 2022) | 1. Activation of L2/3 or L5 PNs<br>2. Simultaneous activation of L2/3 and L5 PNs | Spiking activity averaged across 5050 L5 PNs |

The exact neural elements activated by TMS in different layers of the cortex are unknown. It is hypothesized that TMS at motor threshold activates the axons of L2/3 PNs, which, in turn, monosynaptically activate L5 PNs to produce the I_1_-wave (Di Lazzaro et al. 2012). Moreover, modeling studies suggest large-diameter myelinated axons of L2/3 and L5 PNs have the lowest thresholds for TMS (Aberra et al. 2020, Salvador et al. 2011), and ICMS in NHPs produced robust I-wave responses when the stimulating microelectrode was located in L5 of M1 (Maier et al. 2013, Shimazu et al. 2004). Therefore, we systematically quantified the model response to the activation of PNs from (a) L2/3 alone, (b) L5 alone and (c) L2/3 and L5 together. Long-range afferent inputs from other cortical regions were excluded from the models to isolate the contribution of local intracolumnar circuitry to the short-latency responses. We constructed dose–response curves, where the dose was the proportion of neurons activated, and the response was the spiking activity averaged across both the L5 PNs and PTNs, where available (Table 4). We used the combined L5 PN and PTN axonal spiking activity as the primary output measure across all models to ensure a consistent output metric because not all models contained both neuron types, and we will refer to the combined L5 PN and PTN activity as simply L5 PN activity. This output serves as a proxy for CST axon responses, as these axons are believed to originate from L5 PNs (Lemon 2008). Transmembrane potentials were recorded from L5 PNs axon terminals (Fig. 3A, B), and spike times were obtained from the transmembrane potentials as the time at which crossed −20 mV. The spike times were convolved with a Gaussian function with zero mean and standard deviation of 0.15 ms to account for different conduction speeds (Fig. 3C). Stimulation-pulse-triggered responses were computed for each of the L5 PNs (Fig. 3D), and the responses were averaged across the L5 PNs. The frequency–current curves of L5 PNs across the four models are shown in Extended Data Fig. 3-1. The frequency–current curves varied considerably across models due to differences in biophysical properties of L5 PNs between models (Extended Data Fig. 3-1, 3-2). Dose–response curves were constructed by calculating the number of L5 PNs exhibiting spiking activity at the time interval of an I_1_-, I_2_-, or I_3_-wave.

### Pharmacological modulation

The effects of pharmacological agents on stimulation-induced I-waves are well documented (Di Lazzaro et al. 2012, Lemon 2008, Ziemann 2004, Ziemann & Rothwell 2000). For example, positive allosteric modulators such as benzodiazepines increase the affinity of GABA_A_Rs for GABA (Gielen et al. 2012) and suppress the late phases of TMS-induced I-waves (Burke et al. 1993, Di Lazzaro et al. 2000). Similarly, administration of muscimol, a full agonist for GABA_A_Rs, into the M1 region of the NHP has shown complete suppression of the late I-waves (I_2_ and I_3_) and partial suppression of the I_1_ wave (Johnston 2014, Shimazu et al. 2004). We simulated the effects of positive allosteric modulators on I-waves by increasing the peak conductance (*g*_gaba_a_) of all GABA_A_Rs in the model by 2–4 times the baseline values.

Another prominent effect is the increased magnitude of I-waves during elevated levels of cortical excitability associated with voluntary contraction of contralateral muscles (Di Lazzaro et al. 1998). We simulated the effects of increased cortical excitability on the magnitude of I-waves by increasing the peak conductance of all AMPARs (*g*_ampa_) by 2–4 times the baseline values. We also simulated the effects of an increase in cortical excitability on the magnitude of I-waves by quantifying the model response with and without baseline firing rates. The baseline firing activity was generated using independent Poisson spike trains with uniformly distributed firing rates and delivered to the model neurons through excitatory synaptic connections.

### Virtual lesioning

We dissected the neural circuits responsible for generating I-waves in the models by making virtual lesions of specific synaptic pathways. This was done by setting peak conductance () of the relevant synaptic connections to zero. We computed connection strength matrices across the four models to compare the correlation between connections implicated in the virtual lesion simulations and the strongest excitatory input received by L5 PN from other PNs. Connection strength, as described by Dura-Bernal et al., 2023, was computed using the following equations,

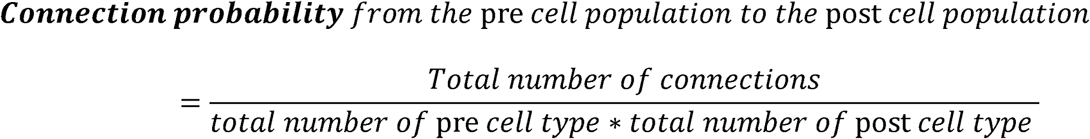

### Numerical simulation

Simulations were implemented in NEURON 7.4 or NetPyNE, with equations solved using the backward Euler method with a time step of 0.025 ms (Hines & Carnevale 2001) and a total simulated time of 20 s. The network simulations were parallelized by dividing the total number of neurons across several CPUs in a round-robin fashion and by setting up the communication of spike times between processors through the Message Passing Interface protocol (Hines & Carnevale 2008). The computational costs for the M1-SC, M1-MC, A1-MC and S1-MC models were approximately 0.5, 40, 96 and 4608 CPU-hours per simulation, respectively.

### Code accessibility

The code to replicate results across the four models will be available on NEURON ModelDB (Ascension #) after publication.

## RESULTS

We used four published models of interconnected networks of cortical neurons, representing different cortical regions, to quantify the short-latency neural responses to stimulation (Table 1). Further, we simulated the effects of various pharmacological agents that alter cortical excitability on the stimulation evoked response. Finally, we simulated selective circuit lesions by disconnecting specific synaptic pathways to understand the microcircuit basis of the neural response.

### L5 pyramidal cell model responses to different stimulation intensities

The exact layer-specific neuron types directly activated by TMS (at motor threshold and higher stimulation intensities) are not fully known. Single-cell modeling studies suggest that axons of PNs have the lowest threshold for activation by TMS (Aberra et al. 2020, Salvador et al. 2011). Similarly, modeling and experimental studies demonstrated that ICMS activates a focal volume of axons around the stimulating microelectrode, comprising axons from PNs across different layers (Histed et al. 2009, Kumaravelu et al. 2022). Further, I_1_ is thought to be generated by activation of the monosynaptic L2/3 PN → L5 PN connection since this connection is one of the strongest in M1 (Di Lazzaro et al. 2012, Douglas & Martin 2004). Therefore, we systematically quantified the spiking activity of L5 PNs in response to activation of (1) L2/3 PNs, (2) L5 PNs, and (3) both L2/3 and L5 PNs. The activation of L2/3 PNs generated I-wave activity in both M1 and A1 models (Fig. 4A, B, C). Although the S1-MC model exhibited rhythmic spiking activity after activation of L2/3 PNs, the timing between peaks (∼5 ms) was substantially longer than the inter-peak interval in experimental I-wave responses (Fig. 4D).

**Figure 4:**
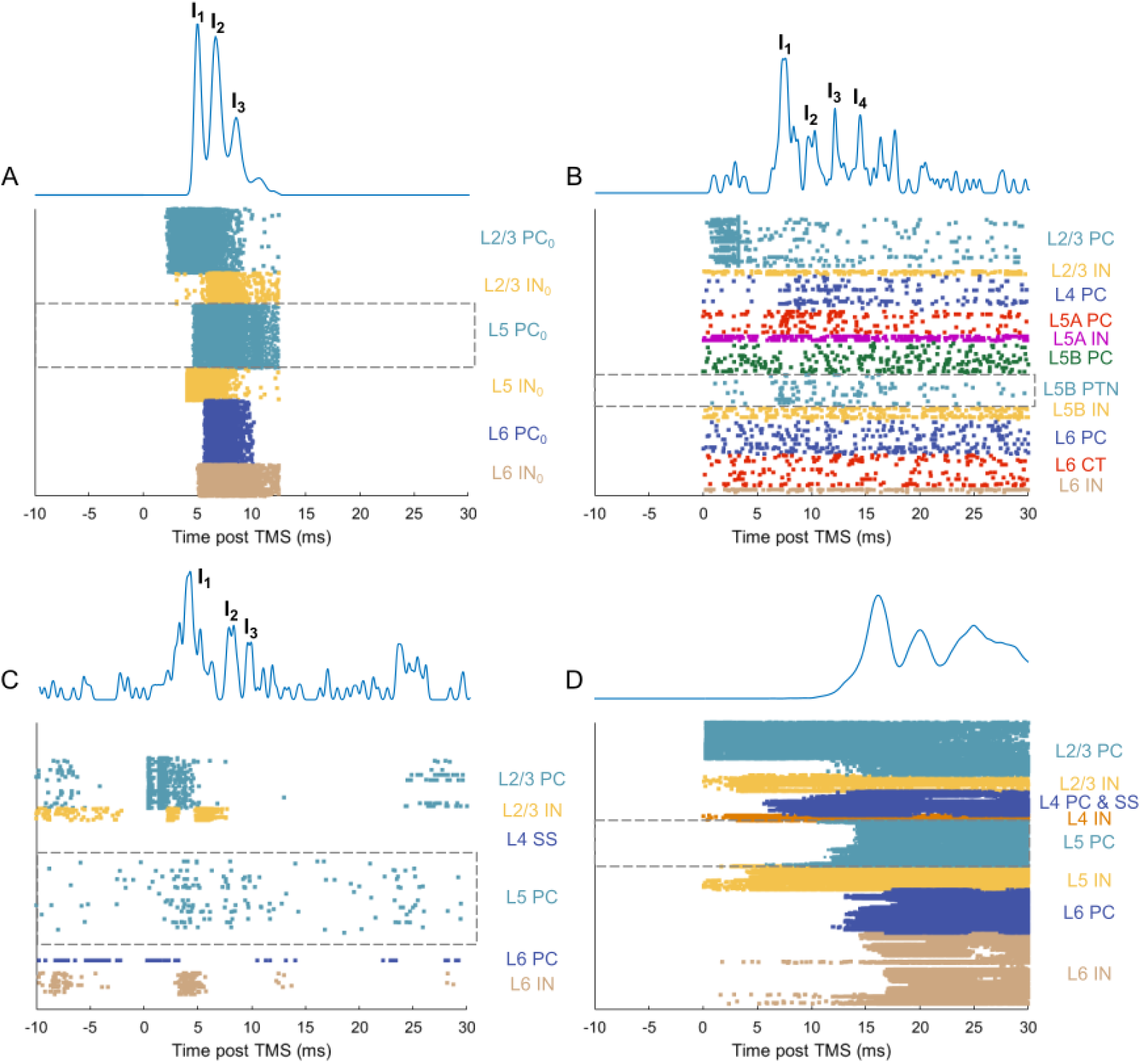
Response of model neurons across different cortical layers to activation of L2/3 PN. Rastergram of spike times of neurons from different layers in the (A) M1-SC model (dose: 40% activation of L2/3 PNs), (B) M1-MC model (dose: 100% activation of L2/3 PNs), (C) A1-MC model (dose: 60% activation of L2/3 PNs) and (D) S1-MC model (dose: 68% activation of L2/3 PNs). The trace at the top of each rastergram represents the smoothed response of all L5 PNs obtained by convolving their firing rates with a Gaussian function. ‘I’ on the traces indicates the I-waves.

The evoked spiking activity of L5 PNs across models was shaped by the interaction of neuronal firing between different layers (Fig. 4). The INs in the deep layers tended to fire in synchrony with the late I-waves, especially I_2_ and I_3_, offering a potential explanation for the suppression seen in the magnitude of late I-waves by GABA_A_R agonists in experimental studies (Fig. 4A, B, C). Further, the magnitude of late I-waves (I_2_, I_3_ and I_4_) became more prominent at higher activation levels compared to lower (Fig. 5 A, B, C). Similarly, rhythmic spiking activity in the S1-MC model was more prominent at higher intensities (Extended Data Fig. 5-1). Responses corresponding to low, medium, and high activation levels are presented for each model. However, the intensity associated with these three activation levels varied between the models. For example, 60% activation of L2/3 PNs in the A1-MC was sufficient to generate a complete I-wave response. In contrast, the M1-MC model required activation of all L2/3 PNs to generate the I-wave response. This discrepancy occurs because each model has unique biophysical properties and synaptic connectivity. For example, the frequency–current curves of L5 PN showed considerable variation across the models (Extended Data Fig. 3-1).

**Figure 5:**
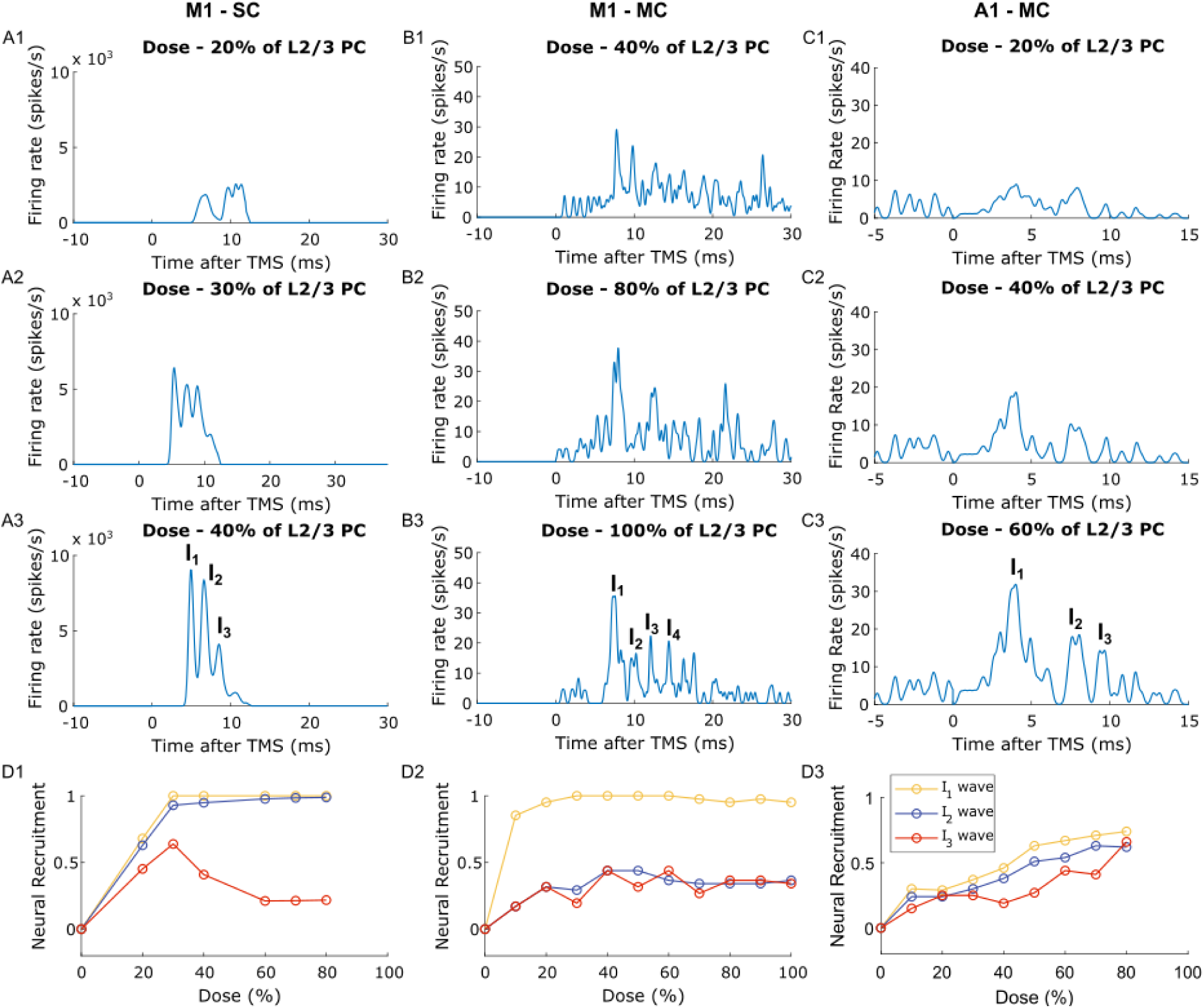
Response of L5 PNs, obtained by convolving their firing rates with a Gaussian function, to activation of different proportions of L2/3 PNs in the (A) M1-SC model, (B) M1-MC model, and (C) A1-MC model. I_1_-I_4_ represent I-waves due to trans-synaptic activation of L5 PNs. The M1-SC, M1-MC, and A1-MC models generated I-waves with response timing closely matching those seen in experimental studies. Number of L5 PNs exhibiting I_1_, I_2_, and I_3_-waves as a function of stimulation intensity in the (D1) M1-SC, (D2) M1-MC, and (D3) A1-MC. The maximum inter-peak interval to detect I-waves was 5 ms. Across the models, the later I-waves (I_2_-I_3_) became more prominent at the higher intensities than the lower intensities, consistent with experimental observations.

We quantified the dose–response curves across models by calculating the number of L5 PNs exhibiting spiking activity at the time intervals of the I_1_, I_2_, and I_3_ waves. At any given activation level, more neurons exhibited the I_1_-wave, followed by the late I-waves (Fig. 5D). In the M1 models, the number of neurons exhibiting I-waves increased with intensity up to a certain dose, followed by saturation at the higher doses (Fig. 5 D1, D2). In the M1-SC model, the number of neurons exhibiting the I_3_ wave increased with stimulation intensity up to a dose of 30% followed by a slight decrease between intensities of 30–60% (Fig. 5D1). At doses > 60%, the number of neurons exhibiting the I_3_ wave saturated (Fig. 5D1). In the A1-MC model, the number of neurons exhibiting I-waves increased with stimulation intensity (Fig. 5D3).

Monophasic TMS in the posterior–anterior direction generates a D-wave in addition to the I-waves at higher stimulation intensities (Di Lazzaro et al. 1998, Di Lazzaro et al. 2012) . Further, ICMS delivered via a microelectrode located in L5 of M1 elicits a D-wave response followed by I-waves (Lemon 2008, Maier et al. 2013, Shimazu et al. 2004). To assess the ability of the models to generate I-waves in conjunction with a D-wave, we quantified the response to direct activation of L5 PN (Extended Data Fig. 4-1). The M1-SC model generated a D-wave followed by three I-waves (I_1_, I_2_ and I_3_), whereas the D-wave in the M1-MC model was followed by only a single I-wave (Extended Data Fig. 4-1 A, B). Although the A1-MC model generated I-waves following the D-wave, the magnitude of the I-waves was greatly diminished compared to the D-wave (Extended Data Fig. 4-1 C). The S1-MC model generated a D-wave followed by I-waves, but the magnitude of this response was substantially smaller compared to the rhythmic spiking activity evoked at longer latencies (∼ 13 ms) (Extended Data Fig. 4-1 D).

Finally, in a subset of simulations, we simultaneously activated PNs from both L2/3 and L5 and measured the response of L5 PNs (Extended Data Fig. 4-2). The M1-SC, A1-MC, and S1-MC models generated a D-wave followed by I-waves (Extended Data Fig. 4-2 A, C, D), while the M1-MC model generated a single I-wave following the D-wave (Extended Data Fig. 4-2 B). These results suggest that a combination of L2/3 PNs and L5 PNs activation, mimicking neural activation with realistic TMS- or ICMS-induced E-fields, can evoke a D-wave followed by I-waves, as seen in the experimental response.

### Modulation of I-waves

We simulated the effects on I-waves of two different interventions – enhanced excitability of cortex, for example from voluntary contraction, and enhanced cortical inhibition from administration of benzodiazepines (Ziemann 2004). The number and amplitude of I-waves increase during voluntary contraction compared to rest (Di Lazzaro et al. 1998). Increasing the peak conductance of AMPARs, intended to represent the increase in M1 excitability resulting from voluntary muscle contraction, enhanced the magnitude of I-waves compared to the control response across all models that generated an I-wave response (Fig. 6A, B, C). The area under the curve for both early and late I-waves increased with higher AMPAR conductance across all models (Fig. 6D). Although the S1-MC model did not generate I-waves at the timing of the experimental I-waves, the model responses to AMPAR modulation were similar to those seen in the other models (Extended Data Fig. 6-1).

**Figure 6:**
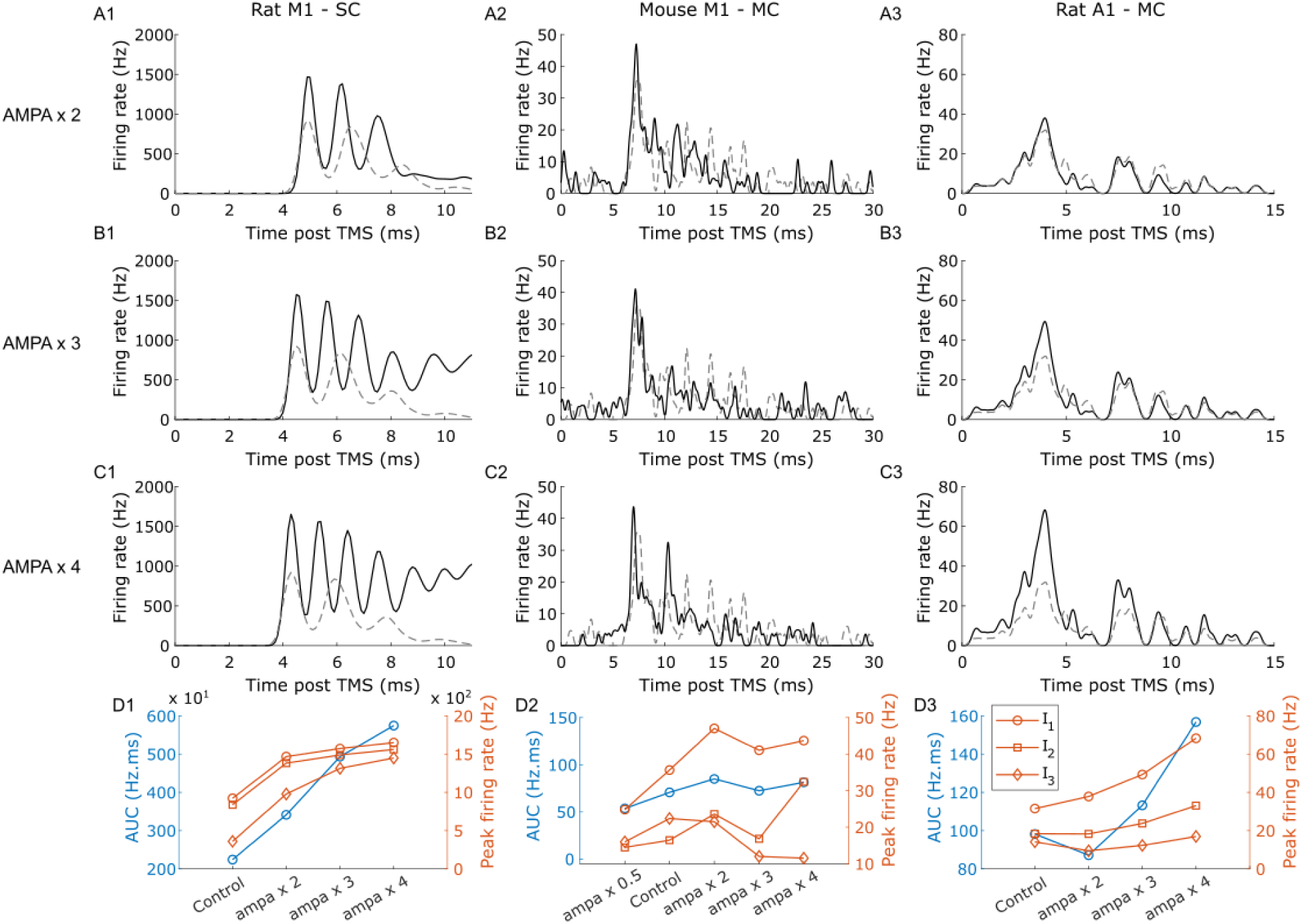
Modulation of model L5 PN excitability mimicking pharmacological modulation of human I-wave responses. Response of L5 PNs, obtained by convolving their firing rates with a Gaussian function, at different levels of AMPA synaptic conductance in the (A) M1-SC model, (B) M1-MC model, (C) A1-MC model. Note that the magnitude of I-waves became more prominent during increased AMPA synaptic conductance (black trace) compared to baseline (dashed gray trace). 60% of L2/3 PNs were activated in the A1-MC model, 100% of L2/3 PNs in the M1-MC model and 40% of L2/3 PNs in the M1-SC model. The area under the response curve (AUC) and the peak response of individual I-waves as a function of AMPA conductance for (D1) M1-SC model, (D2) M1-MC model, (D3) A1-MC model. Across all models, the magnitude of both early and late I-waves increased with AMPA conductance.

An alternative approach to model the increase in cortical excitability from voluntary contraction is enhanced intrinsic firing activity. In contrast to experimental findings during voluntary contraction (Di Lazzaro et al. 1998), the magnitude of the I-waves was substantially decreased in the presence of intrinsic activity compared to response without baseline firing (Extended Data Fig. 6-2). However, this effect was less pronounced for the M1-MC and the A1-MC models (Extended Data Fig. 6-2 B, C). Increasing AMPAR conductance and increasing spontaneous firing rates produced differential effects on the I-wave response. Increasing AMPAR conductance may enhance the effect of stimulation-evoked excitatory synaptic inputs, whereas increasing spontaneous firing may not necessarily increase responsiveness to stimulation-evoked inputs and could increase the proportion of neurons in a refractory state at the time of stimulation. However, the limited information on endogenous activity in cortical neurons during voluntary contraction (such as firing rate, firing patterns, synchrony, etc.) makes modeling these state-dependent effects challenging.

Increasing the peak conductance of GABA_A_Rs synapses, intended to mimic the effect of GABA_A_R receptor positive allosteric modulators, diminished the magnitude of I-waves compared to the control response (Fig. 7A, B, C). The area under the curve of the late I-waves decreased as the GABA_A_R conductance increased in both M1 models (Fig. 7D1, D2), and in the A1-MC model, the magnitude of all I-waves was suppressed compared to the control response (Fig. 7D3). Thus, the models replicated well the experimental effects of these manipulations, strengthening the validity of using the models to decipher the mechanistic origins of I-waves.

**Figure 7:**
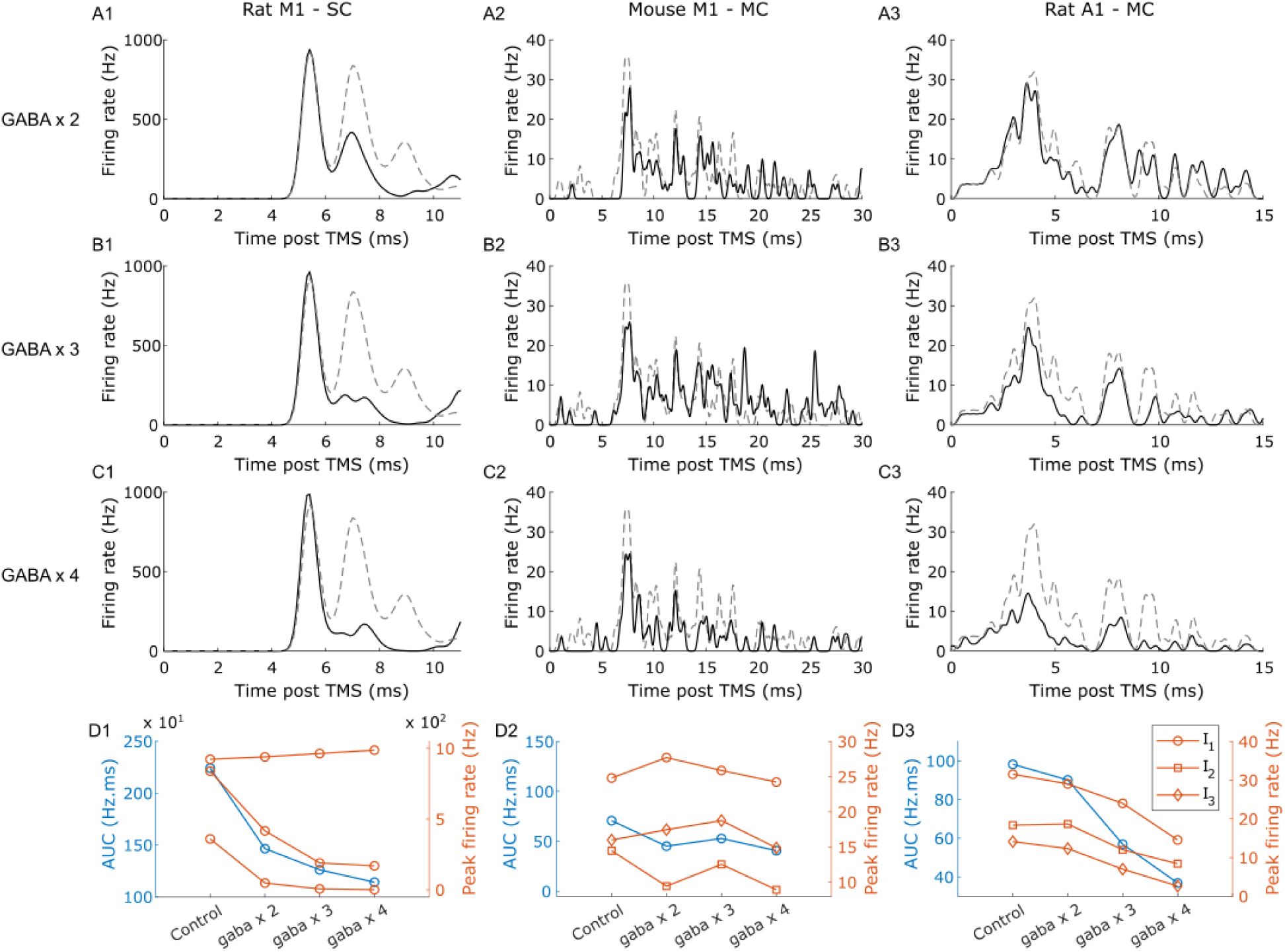
Modulation of model L5 PN excitability mimicking pharmacological modulation of human I-wave responses. Response of L5 PNs, obtained by convolving their firing rates with a Gaussian function, at different levels of GABA-A synaptic conductance in the (A) M1-SC model, (B) M1-MC model, (C) A1-MC model. Note the suppression in magnitude of I-waves during increased levels of GABA-A conductance (black trace) compared to baseline (dashed gray trace). 60% of L2/3 PNs were activated in the A1-MC model, 100% of L2/3 PNs in the M1-MC model and 40% of L2/3 PNs in the M1-SC model. AUC and peak response of individual I-waves as a function of GABA-A conductance for (D1) M1-SC model, (D2) M1-MC model, (D3) A1-MC model. In the M1 models, the magnitude of only the late I-waves decreased with increased GABA-A conductance. The magnitude of both early and late I-waves declined with increased GABA-A synaptic conductance in the A1 model.

### Virtual lesions to dissect the neural origin of I-waves

In the M1 and A1 models, virtual lesioning of the L2/3 PN → L5 PN monosynaptic connection completely silenced the L5 PN firing response to stimulation (Fig. 8A1, D1, B1, D2, C1, D3). Further, removing the recurrent synaptic connection between L5 PNs suppressed the magnitude of the late I-waves in the M1-SC and A1-MC models but not in the M1-MC model (Fig. 8A2, D1, B2, D2, C2, D3). Removing the L6 PN → L5 PN monosynaptic connection did not affect appreciably the I-wave activity compared to the control response (Fig. 8A3, D1, C3, D3). The L5B PTNs in the M1-MC model received excitatory inputs only from PNs in L2/3, L5A, and L5B, other than the recurrent connection from other PTNs in L5B (Fig. 2A1). Since the recurrent connection between L5B PTNs had a very limited effect on the I-waves, synaptic inputs from L5A and L5B PNs generated the late I-waves in the M1-MC model. The results across the two M1 models suggest that the monosynaptic connection from L2/3 PN to L5 PN can contribute to the generation of the I_1_-wave. In contrast, the local connections between the L5 PN and other PNs from L5A and L5B might dominate generation of later I-waves. Next, to determine whether the synaptic pathways implicated in I-wave generation by the virtual lesion simulations corresponded to the strongest synaptic inputs onto L5 PNs, we compared the connectivity matrices across models with the results of the virtual lesion simulations (Fig. 9 and Extended Data Fig. 9-1, 9-2). The strongest monosynaptic connection received by L5 PN from PNs in other layers mirrored the synaptic connections implicated in the generation of the I-waves through the virtual lesion simulations (Fig. 9). For example, in the M1-MC model, L2/3 PN to L5 PN was the strongest incoming connection to L5 PN followed by the recurrent connection from L5 PN (Fig. 9B), and these two connections were implicated in the generation of the I-waves in the virtual lesion simulations. This correspondence between the connections implicated in the virtual lesion simulations and the strongest connection received by L5 PN was also present across the other models (Fig. 9 and Extended Data Fig. 9-1, 9-2).

**Figure 8:**
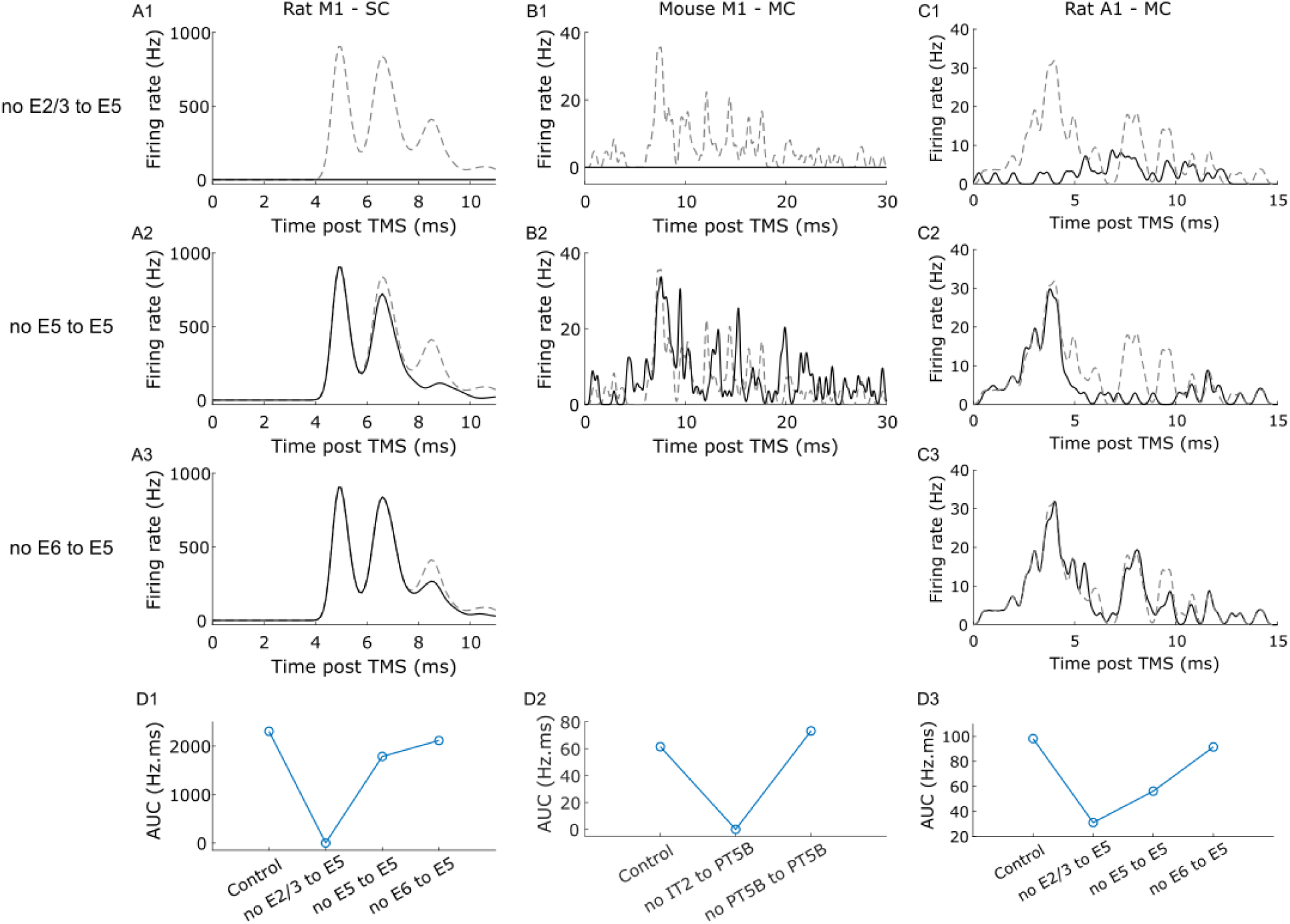
Dissection of the cortical circuit element contributions to stimulation-induced I-waves by disconnecting individual synaptic connections in the cortical column. Response in L5 PNs, obtained by convolving their firing rates with a Gaussian function, after virtual lesioning of individual synaptic connections in the (A) M1-SC model, (B) M1-MC model, (C) A1-MC model. The activation in each model was: A1-MC model – 60% activation of L2/3 PNs, M1-MC model – 100% activation of L2/3 PNs and M1-SC model – 40% activation of L2/3 PNs. In panels (A1-A2, B1-B2, C1-C2) note the suppression of I-waves (black trace) compared to control (dashed gra trace). Lesioning neurons in other layers did not substantially affect the I-wave activity. AUC after virtual lesioning of individual synaptic connections in the (D1) M1-SC model, (D2) M1-MC model, (D3) A1-MC model.

**Figure 9:**
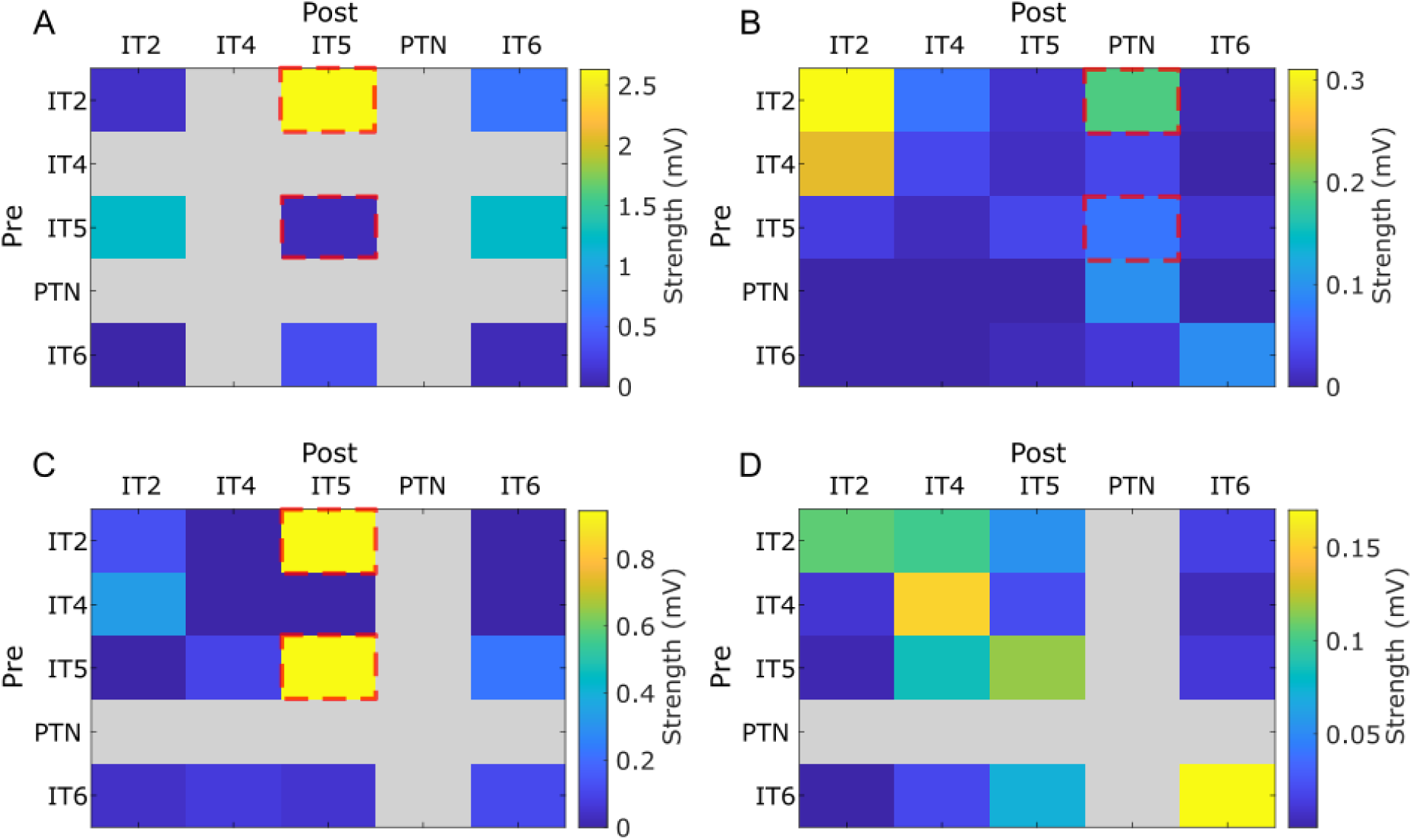
Strength of monosynaptic excitatory connections in (A) M1-SC model, (B) M1-MC model, (C) A1-MC model, (D) S1-MC model. Connection strength was computed using the equation, . The red dashed box indicates the synaptic connections implicated in the generation of I-waves in the virtual lesion simulations. Gray boxes indicate synaptic connections that do not exist in the model. Since the S1-MC did not generate rhythmic spiking activity at the timing of the experimentally observed I-waves, no virtual lesion simulations were performed in that model (hence no red dashed box).

### L5 PN recruitment dynamics across cortical models

To investigate why the S1-MC model generated rhythmic spiking activity with longer inter-peak intervals than the responses observed in the other models, we quantified the contribution of individual L5 PNs to each population volley within the rhythmic firing response (Fig. 10). We identified two distinct patterns of L5 PN recruitment across the four models. First, in the M1-MC and A1-MC models, L5 PNs that participated in the first volley (i.e., the I_1_ response) showed little to no participation in the I_2_ response (Fig. 10B), indicating that later I-wave responses were generated predominantly by different L5 PNs. In contrast, in the M1-SC and S1-MC models, a large number of L5PNs (> 90%) activated during the I_1_ response also fired during the second volley (Fig. 10B), indicating substantial re-recruitment of the same L5 PNs. However, the re-recruited L5 PNs in the S1-MC model showed a longer recovery period before the next spike (∼3 ms) than in the M1-SC model (∼2 ms). This longer re-firing timescale of L5 PNs in the S1-MC model was consistent with the longer inter-peak intervals between its population volleys compared with the other models (Fig. 10C).

**Figure 10:**
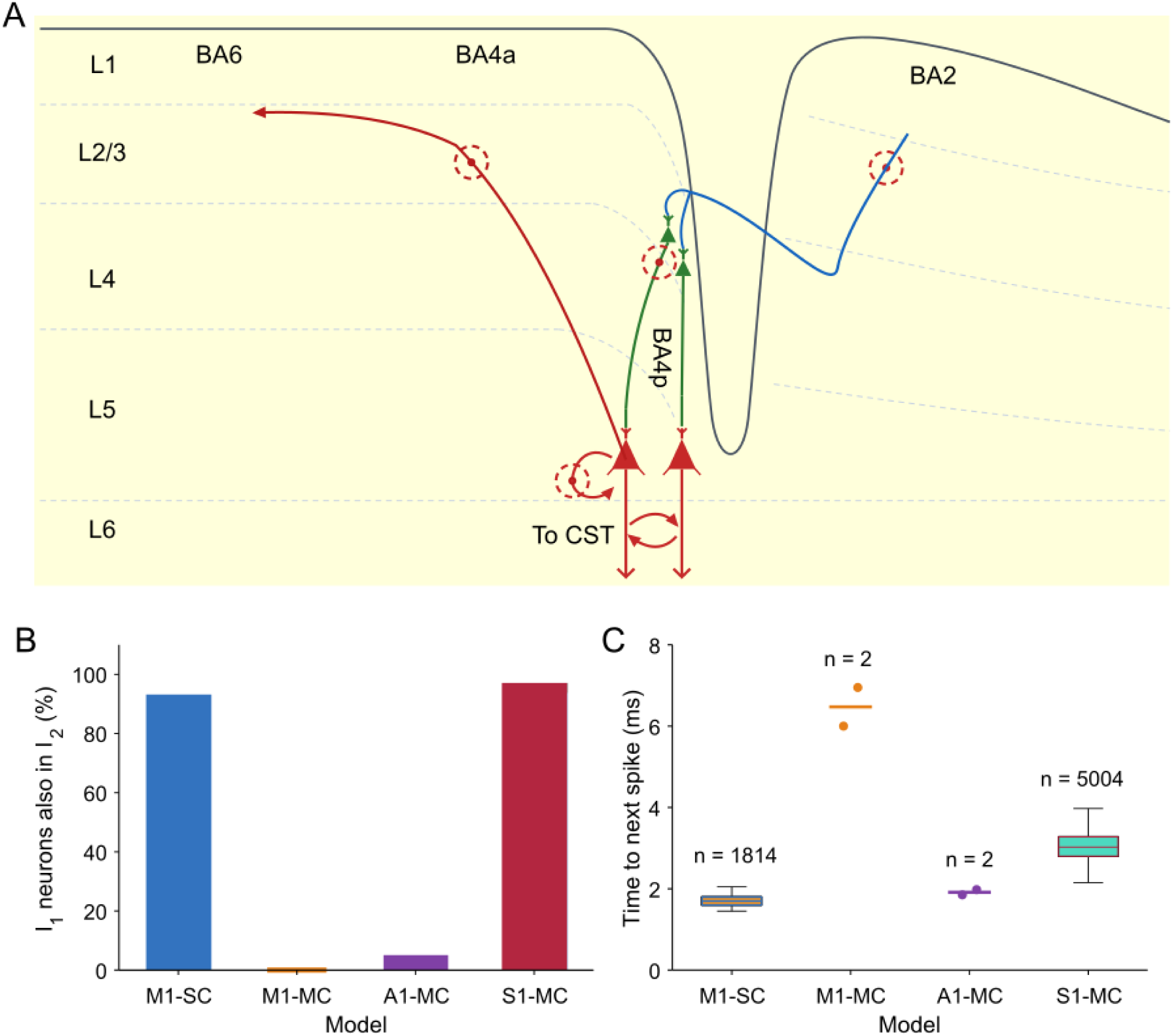
Mechanism of I-wave generation. (A) Schematic of cortical circuitry implicated in I-wave generation following cortical stimulation. Descending CST output arises from L5 PNs in Brodmann area (BA) 4p, which receive direct and trans-synaptic inputs from local and cortico-cortical pathways. Because this study aimed to isolate intracolumnar effects on I-wave generation, we excluded long-range inputs into BA4p from other cortical regions. The red dashed circles indicate axons that can be directly activated by cortical stimulation modalities such as TMS and ICMS. While TMS can induce broadband axonal activation (i.e., both long-range and local), only axons of local L2/3 and L5 PNs within BA4p are expected to be activated during ICMS. ICMS of BA4p has been shown to be sufficient to generate I-waves. The virtual lesion simulations implicated local L2/3→L5 and L5→L5 synaptic pathways within BA4p in I-wave generation. (B) Percentage of L5 PNs participating in the first population volley that also fired in the second population volley across the cortical models. M1-SC and S1-MC exhibited substantial re-recruitment of the same L5 PN population in the consecutive volleys, whereas little or no overlap was observed in the M1-MC and A1-MC models. This implies different neurons contributed to each I-wave volley in the M1-MC and A1-MC models, whereas in the M1-SC and S1-MC models the same neurons contributed to the rhythmic spiking activity. (C) Time from the spike of an L5 PN during the first population volley to the next spike of the same neuron within the subsequent three population volleys. M1-SC exhibited rapid re-firing of the L5 PNs that participated in the first volley, whereas L5 PNs in the S1-MC model exhibited a longer re-firing interval. This is why L5 PN rhythmic activity in the S1-MC model showed a longer inter-peak interval (∼5 ms) than in the other models and mismatched with the timing of the experimentally observed I-waves. n denotes the number of first-volley L5 PNs that refired within the subsequent three volleys. Box plots indicate the median and interquartile range, with whiskers indicating the outliers.

## DISCUSSION

The microcircuit basis for complex responses to TMS and ICMS is unclear, and we implemented four published network models, representing different cortical regions, to study the short-latency neural responses to stimulation. We quantified the responses to different proportions of activation of PNs from either L2/3, L5, or both, as well as studied the modulation of the responses to interventions that altered excitability. Both M1 and A1 models generated I-waves following activation of PNs in L2/3 and L5. Although the S1 model exhibited rhythmic spiking activity after activation of L2/3 PNs, the timing between peaks did not match experimental I-waves. Increasing the strength of GABA_A_R synapses in the models, mimicking the effect of benzodiazepines and muscimol, decreased the magnitude of the late I-waves compared to the control response. Virtual lesion simulations revealed that in these models the I_1_-wave was generated by the L2/3 to L5 PN connection, while the recurrent connections between L5 PNs generated the later I-waves.

### Comparison of model and experimental responses to TMS and ICMS

There is only a single study reporting the short-latency response to TMS recorded from L5 PNs in rodent M1 (Li et al. 2017). The experimental response to monophasic TMS in the posterior–anterior direction was obscured by stimulation artifact for ∼1 ms after the TMS pulse, and the response beyond 1 ms comprised two short-latency peaks at 1.25 ms and 3.75 ms. Further, the response became more prominent at higher intensities. The model responses included short-latency peaks similar to these experimental recordings. The magnitude of the response components in the model became more prominent at the higher intensities, as seen in the experimental recordings. Although the responses across models qualitatively matched well with the experimental data, the limitations associated with the experimental data make it challenging to compare quantitatively the responses between the model and experiment. First, there is a paucity of single-unit responses from cortical neurons in the literature, especially in the short-latency time scale (1–10 ms post-TMS). Second, the Li et al. study shows only multi-unit spiking responses averaged across four rats, making it infeasible to validate model-based single-unit responses against experimental data. Finally, Li et al. reports the response to TMS normalized by the baseline firing rate of L5 PN but did not report the baseline firing rate of the L5 PN. Since our modeling results show that the response evoked by stimulation is sensitive to the baseline firing rate of the L5 PN, making a direct comparison between the model responses and experimental recordings is challenging. Although no studies have reported the short-latency L5 PN response to ICMS in M1, one study quantified the PN response to ICMS in S1 of NHP (Sombeck et al. 2022). The PN response to S1 stimulation exhibited rhythmic spiking activity, akin to D- and I-waves, within <10 ms, similar to the L5 PN response observed upon activation of L5 PN in the S1-MC model (Extended Data Fig. 4-2 D).

There are no studies directly quantifying the effect of AMPA receptor modulation on the I-wave response. The thresholds for eliciting late I-waves decreased during voluntary contraction compared with the rest condition (Di Lazzaro et al. 1998). Although the threshold to evoke an I_1_-wave was reduced during voluntary contraction compared to rest in 1/3 patients, the effect was unclear in the remaining two patients (Di Lazzaro et al. 1998). Further, the amplitude of individual I-waves increased during voluntary contraction compared to rest (Di Lazzaro et al. 1998). The magnitude of I-waves (I_1_, I_2_, and I_3_) increased with AMPAR conductance across models, consistent with the enhanced I-wave responses observed during voluntary contraction and suggesting a potential role for increased excitability and agreeing well with the limited experimental results. In an experimental study quantifying the effect of the GABA_A_R positive allosteric modulator, the magnitude of the late I-wave evoked by TMS decreased, whereas the I_1_-wave magnitude was unaffected (Di Lazzaro et al. 2000). In both M1 models, the magnitude of the I_1_-wave was unaffected by an increase in GABA_A_R conductance, while the magnitude of the late I-waves decreased with increased GABA_A_R synaptic strength. However, the magnitude of all three I-waves decreased with an increase in GABA_A_R conductance in the A1 model. In contrast to TMS, an ICMS study in M1 (L5) of NHP showed that the magnitude of the I_1_-wave decreases in the presence of the GABA_A_R agonist muscimol, whereas the late I-waves were completely suppressed and the D-wave remained mostly unaffected (Shimazu et al. 2004). There are two possible mechanisms by which GABA_A_R modulators can cause a selective suppression of late I-waves without affecting the I_1_-wave. First, the inhibitory interneurons could fire in synchrony at the timing of the late I-waves and in combination with the effects of drug, suppress strongly the magnitude of late I-waves. Deep inhibitory interneurons across models fired in synchrony with the I_1_-wave as well as with late I-waves, but there was increased synchrony with the late I-waves compared to the I_1_-wave (Fig. 4). The second mechanism by which the I_1_-wave would remain unaffected by GABA_A_R modulation is when the I_1_-wave is generated as a result of direct activation of long-range axons that project from Brodmann area BA4p to BA4a and would not involve a trans-synaptic mechanism (Fig. 10A).

### Neural origin of I1 responses

There are several proposed mechanisms regarding the neural origin of the I_1_ response and we discuss each of these mechanisms in detail below (Fig. 10A): (1) monosynaptic activation of L2/3 to L5 PN connections, (2) activation of L5 PN recurrent connections, and (3) activation of long-range axon collaterals projecting from Brodmann area BA4p to BA4a.

Regarding the first mechanism, the I_1_-wave is thought to be produced by monosynaptic, excitatory inputs to L5 PNs from L2/3 (Di Lazzaro et al. 2012). This hypothesis is based on two observations. First, there are strong excitatory inputs from L2/3 to L5 that are often emphasized in canonical cortical column models (Anderson et al. 2010, Douglas & Martin 2004, Thomson & Bannister 2003). Second, TMS in the PA direction at motor threshold generates I_1_ responses but not a D-wave, thus ruling out direct activation of L5 PN main axons by threshold TMS. In line with this observation, results from our network models indicate that I_1_ can be generated by the monosynaptic excitatory connection from the L2/3 PN to L5 PN.

Regarding the second mechanism, ICMS of L5 in M1 of NHP elicits a D-wave followed by a series of I-waves (Maier et al. 2013, Shimazu et al. 2004). Further, single-cell modeling studies suggest that L5 PN axon collaterals have the lowest threshold to TMS (Aberra et al. 2020, Salvador et al. 2011). Our modeling results suggest I-waves can be generated following a D-wave after activation of PNs in L5 via recurrent connections between L5 PNs (i.e., collaterals that synapse onto other L5 PNs).

Finally, activation of L5 PN long-range axon collaterals that project from Brodmann area BA4p (M1 in the deep sulcal wall) to Brodmann area BA4a (M1 more towards the superficial gyral crown) could generate a response at the latency of an I_1_-wave (Massimini et al. 2026) (Fig. 10A). The action potentials would travel antidromically along the collaterals and then propagate re-orthodromically down the main L5 PN axons; this propagation delay could result in the I_1_-wave timing without the involvement of synapses. This theory is plausible due to the relatively high thresholds for direct activation of PNs in the BA4p regions that project strongly to the CST because of their location in the deeper sulcal wall. In comparison, the areas close to the gyral crown (BA4a regions) likely have lower thresholds for direct axonal activation due to the higher electric fields from TMS (Aberra et al. 2020, Bungert et al. 2017, Weise et al. 2020). This mechanism is not represented in the models examined here, which do not incorporate the long-range axonal geometry and associated conduction delays required to generate I_1_-wave through antidromic and subsequent re-orthodromic propagation.

Although activation of long-range axon collaterals might be sufficient to generate a response at the timing of the I_1_-wave, our results suggest that excitatory synaptic inputs to L5 PNs from L2/3 PNs and L5 PNs might still contribute to the generation of the I_1_-wave (Fig. 10A). For example, muscimol injection in M1 reduces the I_1_-wave amplitude but does not abolish it (Shimazu et al. 2004). Similarly, cooling of the motor cortex preferentially reduces the late I-waves, whereas removal of the precentral cortex abolishes I-waves evoked by stimulation of cortical regions projecting to M1, supporting trans-synaptic activation of motor cortical PTNs through corticocortical inputs (Amassian et al. 1987). These and other findings suggest that both synaptic and non-synaptic mechanisms could contribute to the I_1_-wave, especially at suprathreshold stimulation (Massimini et al. 2026, Worbs et al. 2026, Ziemann 2020). The primary objective of this study was not to model the direct (i.e., non-synaptic) neuronal activation produced by TMS- or ICMS-induced E-fields, since realistic stimulation-induced fields were not incorporated into the models. Instead, using detailed cortical network models, we investigated the subsequent synaptic mechanisms elicited by this initial neuronal activation.

### Neural origin of late I responses

Three hypotheses exist regarding the origin of later I-waves (I_2,_ I_3…_). First, because the magnitude of late I-waves is suppressed by GABA_A_R positive allosteric modulators (Di Lazzaro et al. 2000), it has been hypothesized that inhibitory interneurons might play a role in generating the late I-wave response. We corroborated this experimental observation through simulations of increased GABA_A_R conductance. Second, L5 PN form reciprocal connections with L2/3 PN (Di Lazzaro et al. 2012), and it is hypothesized that these reciprocal connections between L5→L2/3→L5 PN might generate late I-waves (Di Lazzaro et al. 2012). Regarding the first hypothesis, for GABA_A_R modulators such as benzodiazepines to act, there should be a release of GABA from inhibitory interneurons (Olsen 2018, Usdin 1982). However, inhibitory interneurons need not fire synchronously with the late I-waves, as GABA_A_R-mediated inhibitory conductances persist beyond the initiating interneuron spike and can therefore suppress the magnitude of the late I-waves. Across the models, GABA_A_R conductances had decay time constants ranging from approximately 6 ms to 100 ms, substantially longer than the ∼1.5 ms interval between successive I-waves. Deep inhibitory interneurons across our models exhibited increased synchrony with the late I-waves, while increasing GABA_A_R conductance suppressed the late I-wave magnitude. Together, these results support a modulatory role of inhibitory interneurons in shaping the late I-wave response rather than a direct role in generating the late I-waves .

Concerning the second hypothesis, although the recurrent connections through L5→L2/3→L5 PN generating I-waves is plausible, we ruled out the possibility of this pathway generating the late I-wave response since the connection from L5 to L2/3 PN was very weak across both the M1 models (Fig. 9). Our modeling results indicate that the L5→L5 PN recurrent excitatory connection mediated by AMPA receptors is more likely to generate late I-wave-like activity. The timing of a monosynaptic recurrent excitatory connection from L5 PN to L5 PN mediated by AMPA receptors is expected to be ∼1–1.5 ms (Ghosh & Porter 1988, Morishima & Kawaguchi 2006, Morishima et al. 2011), within the range of the timing of I-waves. Indeed, the recurrent excitatory synaptic connection between L5 PNs was thought to mediate the short-latency response to TMS in M1 in recent experimental studies (Kurz et al. 2019, Li et al. 2017).

Finally, activation of afferent inputs to M1 from other cortical regions during TMS, including the premotor cortex and somatosensory cortex, can contribute to the generation of late I-waves (Massimini et al. 2026, Siebner et al. 2022) (Fig. 10A). For instance, ICMS of the premotor cortex facilitated the generation of I-waves in response to M1 stimulation; however, premotor cortex stimulation alone was insufficient to generate I-waves (Shimazu et al. 2004). We removed the afferent inputs across models since the focus of this study was to isolate intra-columnar effects in I-wave generation.

### Limitations of the model

Our biophysically based models representing different cortical regions are the most advanced models for studying the short-latency responses to TMS and ICMS, but they have several important limitations. First, the cortical column models are based on the anatomy and physiology of the rodent brain, whereas the I-wave response to TMS and ICMS has been chiefly characterized in humans and NHPs. Although very few studies exist quantifying the short-latency response to TMS in rodents, a study by Li et al. reported an I-wave-like response in L5 PNs during TMS over M1, and our model results agreed well with the experimental recordings (Li et al. 2017). However, the model predictions regarding the mechanisms underlying the short-latency response need to be validated experimentally. Second, TMS and ICMS at different pulse amplitudes generate, proportionally, different electric field intensities in the brain. However, the strength of stimulation was not modeled in terms of electric field but instead as levels of activation of PNs. Since the electric field was not explicitly modeled, we had to make assumptions about the neural types activated by different stimulation intensities. Third, the models lacked medium- and long-range axonal projections, as well as their collaterals, between M1 and other cortical regions, which may contribute to the generation of I-waves (Massimini et al. 2026). Although no studies have quantified M1/CST responses during TMS applied to non-motor regions, ICMS in the premotor cortex facilitates the late I-wave responses evoked by M1 stimulation in NHP (Shimazu et al. 2004). Finally, we did not model the long-latency responses to TMS and ICMS. TMS over M1 generated short-latency I-wave responses (1–6 ms) considered here, followed by a long-latency excitatory and inhibitory responses (6–200 ms) and a rebound excitation between 200–300 ms (Li et al. 2017, Romero et al. 2019). ICMS also elicits long-latency responses similar to those seen with TMS (Butovas & Schwarz 2003, Sombeck et al. 2022). However, we did not attempt to model the long-latency responses as some of the models (M1-MC and A1-MC) lacked GABA_B_R mechanisms and feedback connections from thalamus, which are believed to mediate the long-latency inhibitory and post-inhibitory rebound responses to electrical stimulation (Butovas et al. 2006, Kumaravelu & Grill 2024).

## Conclusion

We used detailed cortical network models to investigate the circuit mechanisms underlying the short-latency responses to TMS and ICMS across multiple cortical regions. Our simulations demonstrate that distinct cortical microcircuits can reproduce experimentally observed D- and I-wave responses and provide mechanistic insight into the circuit motifs that generate them. Although the models do not capture every aspect of cortical physiology, they provide a unified computational framework for systematically testing competing mechanistic hypotheses, evaluating circuit contributions through virtual perturbations, and guiding the interpretation of experimentally measured TMS- and ICMS-evoked cortical responses across different regions.

## Author Contributions

K.K., G.J.Y., A.S.A., M.A.S., A.V.P., and W.M.G. conception and design of research; K.K. ran model simulations; K.K. analyzed data; K.K., G.J.Y., A.S.A., M.A.S., A.V.P., and W.M.G. interpreted results of model simulations; K.K. prepared figures and drafted initial version of manuscript; K.K., G.J.Y., A.S.A., M.A.S., A.V.P., and W.M.G. edited and revised manuscript; K.K., G.J.Y., A.S.A., M.A.S., A.V.P., and W.M.G. approved final version of manuscript.

## Acknowledgments

Research reported in this publication was supported by the National Institute of Neurological Disorders and Stroke and the National Institute of General Medical Sciences of the National Institutes of Health under Award Numbers R01NS117405, R01NS088674, and R25GM103765. The content is solely the responsibility of the authors and does not necessarily represent the official views of the National Institutes of Health. We thank the Duke Compute Cluster team for computational support. The authors would like to thank Boshuo Wang and Corrie Camalier for their helpful discussions on this work.

## Conflict of Interest

A.V.P. is an inventor on patents and patent applications on transcranial magnetic stimulation technology and has received patent royalties and consulting fees from Rogue Research; equity options, scientific advisory board membership, and consulting fees from Ampa Health; equity options and consulting fees from Magnetic Tides; consulting fees from Soterix Medical; equipment loans from MagVenture; hardware donation from Magstim; and research funding from Motif. Preliminary results from this work were presented at the 4^th^ International Brain Stimulation Conference (Charleston, SC, USA, Dec. 2021), the Society for Neuroscience’s 52^nd^ Annual Meeting (Washington DC, USA, Nov. 2023), and the 6^th^ International Brain Stimulation Conference (Kobe, Japan, Feb. 2025).

**Glossary of acronyms**
TMS: Transcranial magnetic stimulation
M1: Primary motor cortex
S1: Primary somatosensory cortex
A1: Primary auditory cortex
MC: Multi-compartment
SC: Single compartment
MEP: Motor evoked potential
CST: Corticospinal tract
NHP: Non-human primate
ICMS: Intracortical microstimulation
PN: Pyramidal neuron
FRB: Fast rhythmic bursting
RS: regular spiking
FS: fast spiking
LTS: Low-threshold spiking
PTN: pyramidal tract neuron
CT: Cortico–thalamic
TC: thalamocortical
TCR: thalamocortical relay
nRT: reticular nucleus γ-aminobutyric acid
GABA: α-Amino-3-hydroxy-5-methyl-4-isoxazolepropionic acid
AMPA: N-methyl-D-aspartic acid NMDA
BA: Brodmann area.

