## Extended Data for "Computational investigation of circuit mechanisms underlying short-latency responses to cortical stimulation"

**Supplementary Figures (Extended Data)**

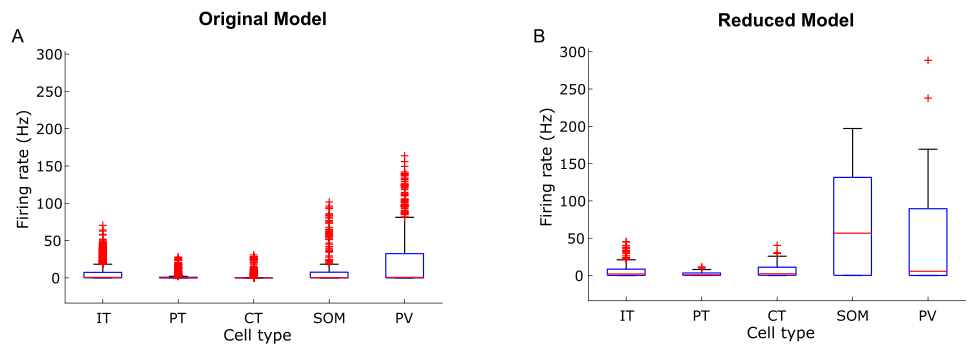

**Figure 2-1:** Comparison of firing rates across cell types between (A) the original M1-MC model and (B) the reduced M1-MC model.

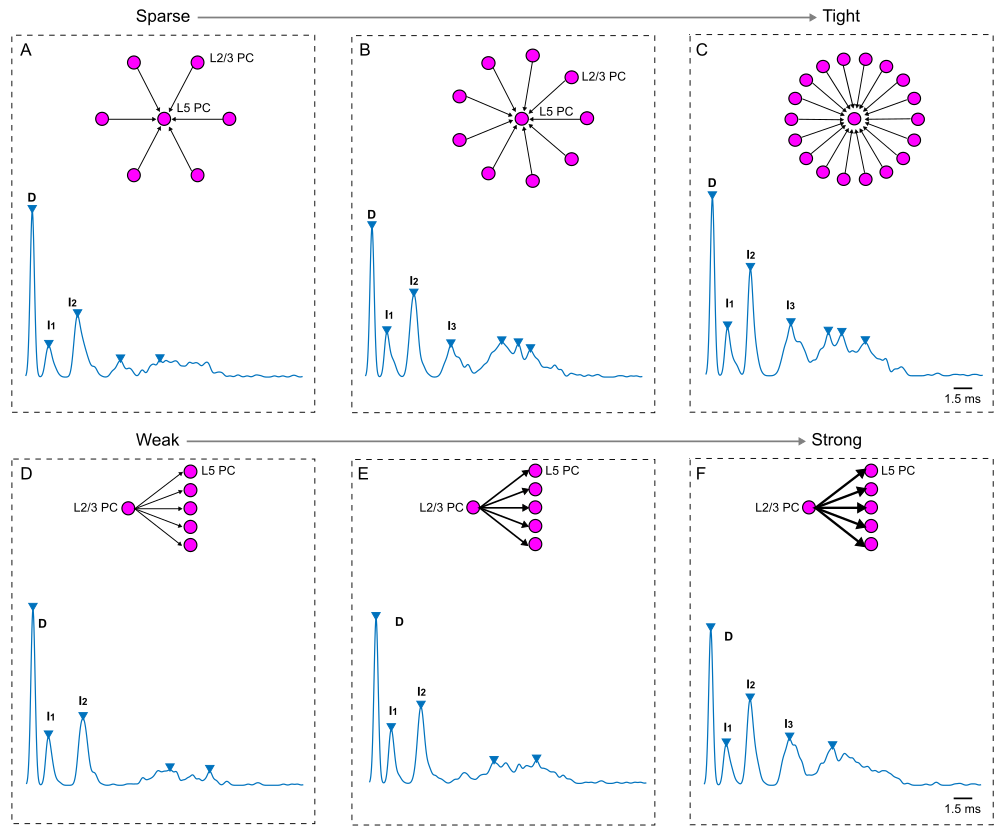

**Figure 2-2:** Influence of L2/3 PN → L5 PN synaptic parameters on the L5 PN spiking response, obtained by convolving their firing rates with a Gaussian function*,* to stimulation in the Traub model. (A-C) L5 PN response at different connection architectures (sparse to tight) between L2/3 PN and L5 PN. (D-F) L5 PN response at various connection strengths (weak to strong) between L2/3 PN and L5 PN.

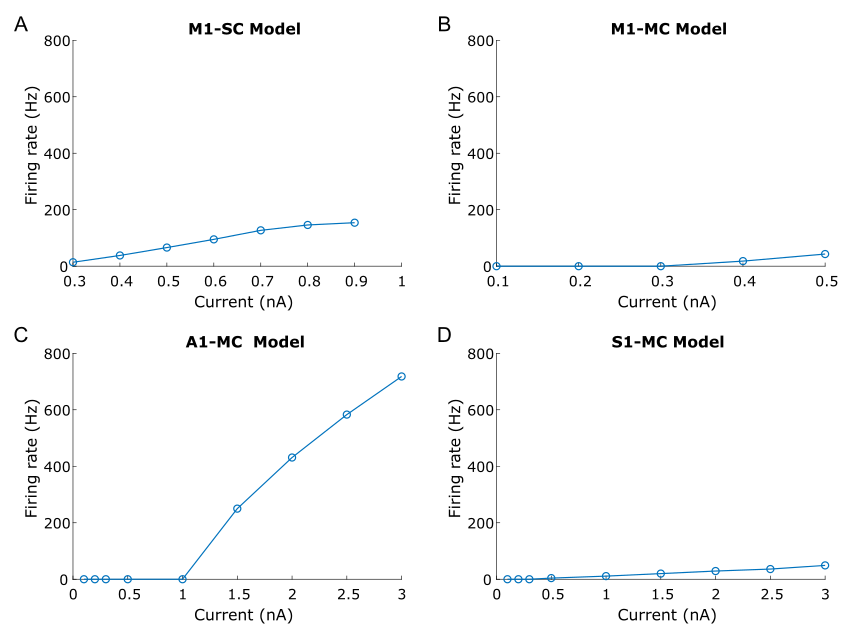

**Figure 3-1:** Firing behavior of L5 PN across the four models used in the study. Frequency–current curves for L5 PN in (A) M1-SC model, (B) M1-MC model, (C) A1-MC model, and (D) S1-MC model. There was a reduction in firing rate of L5 PN in M1-MC model at currents > 0.5 nA (for additional details, please see Extended Data Fig. 3-2).

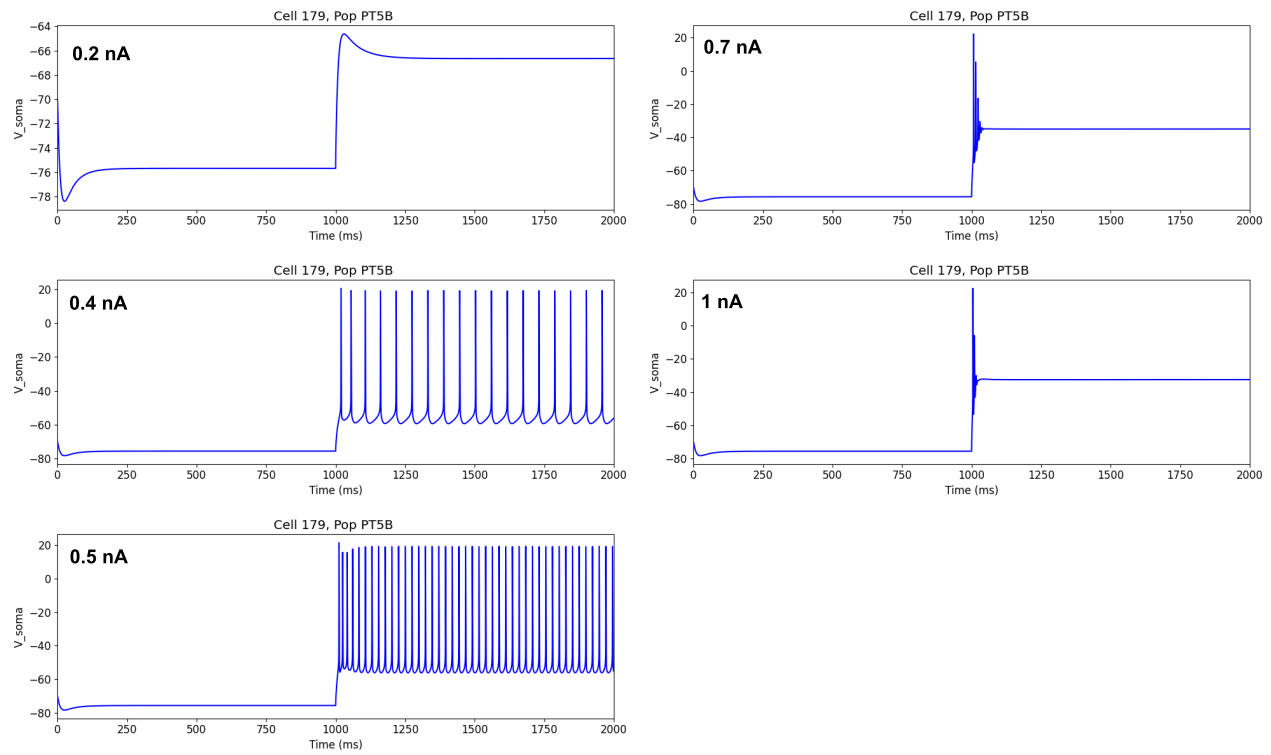

**Figure 3-2:** L5 PN membrane potential recordings to various current levels in the M1-MC model. The firing rate increased with current levels up to 0.5 nA. At higher current levels (>0.7 nA), the model neurons exhibited an initial burst of activity followed by sustained depolarization with no spiking activity.

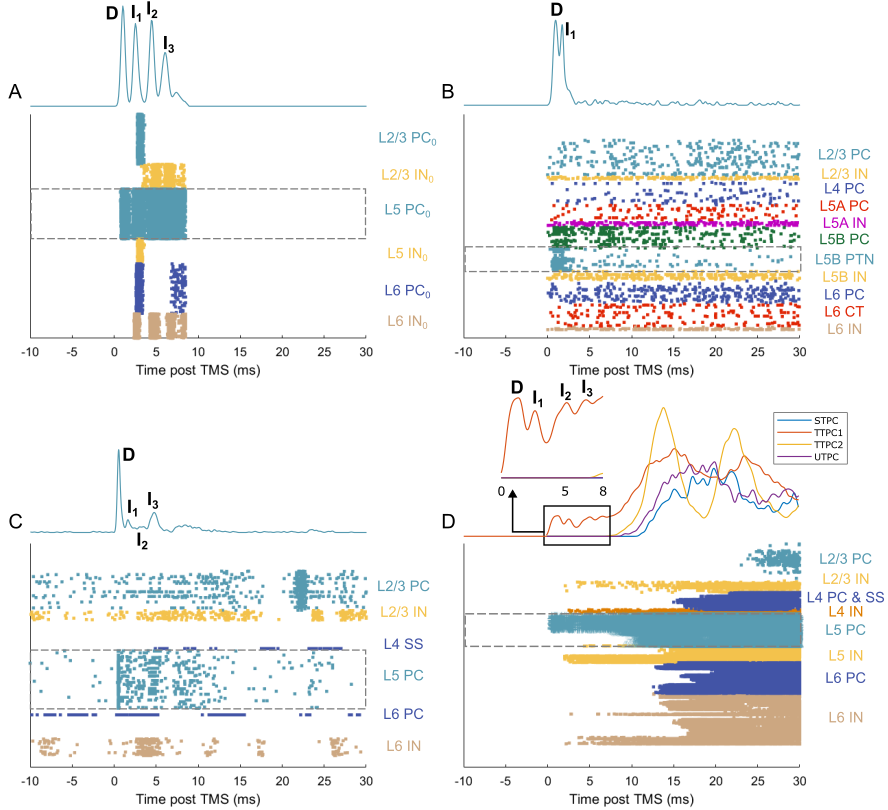

**Figure 4-1:** Response of model neurons across different cortical layers to direct activation of L5 PNs. Rastergram of spike times of neurons from different layers in the (A) M1-SC model (dose: 70% activation of L5 PNs), (B) M1-MC model (dose: 60% activation of L5 PNs), (C) A1-MC model (dose: 60% activation of L5 PNs) and (D) S1-MC model (dose: 60% activation of L5 PNs). The trace at the top of each rastergram represents the smooth response of L5 PNs obtained by convolving their firing rates with a Gaussian function. ‘D’ and ‘I’ on the traces indicate the D- and I-waves.

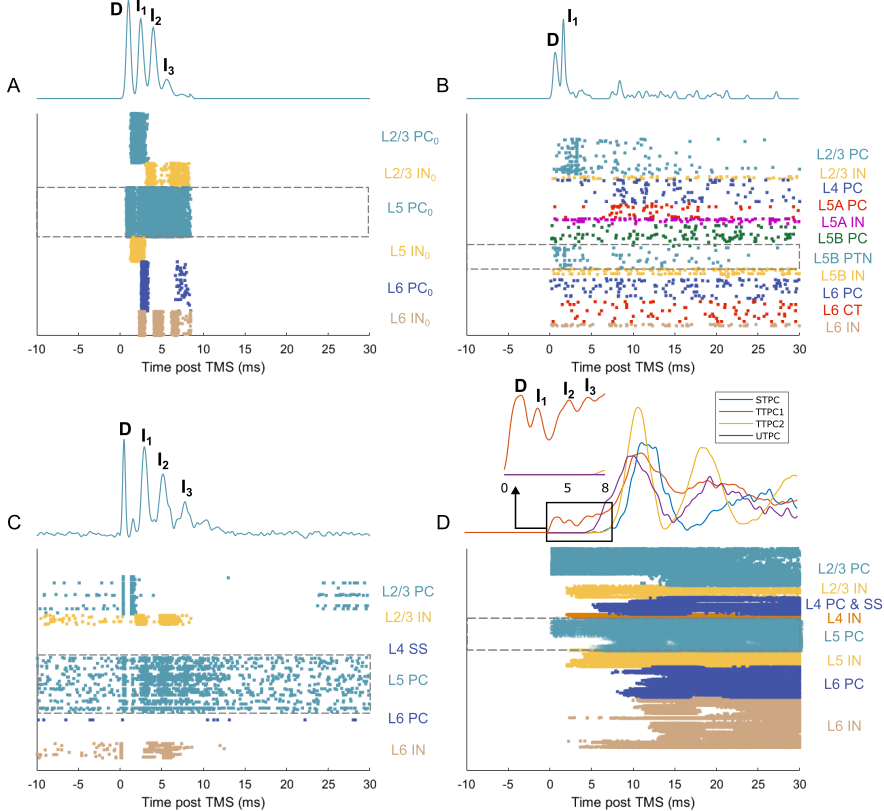

**Figure 4-2:** Response of model neurons across different cortical layers to simultaneous direct activation of L2/3 and L5 PNs. Rastergram of spike times of neurons from different layers in the (A) M1-SC model (dose: 40% activation of L2/3 PNs and 80% activation of L5 PNs), (B) M1-MC model (dose: 80% activation of L2/3 PNs and 40% activation of L5 PNs), (C) A1-MC model (dose: 60% activation of L2/3 PNs and 40% activation of L5 PNs) and (D) S1-MC model (dose: 68% activation of L2/3 PNs and 60% activation of L5 PNs). The trace at the top of each rastergram represents the smooth response of L5 PNs obtained by convolving their firing rates with a Gaussian function. ‘D’ and ‘I’ on the traces indicate the D- and I-waves.

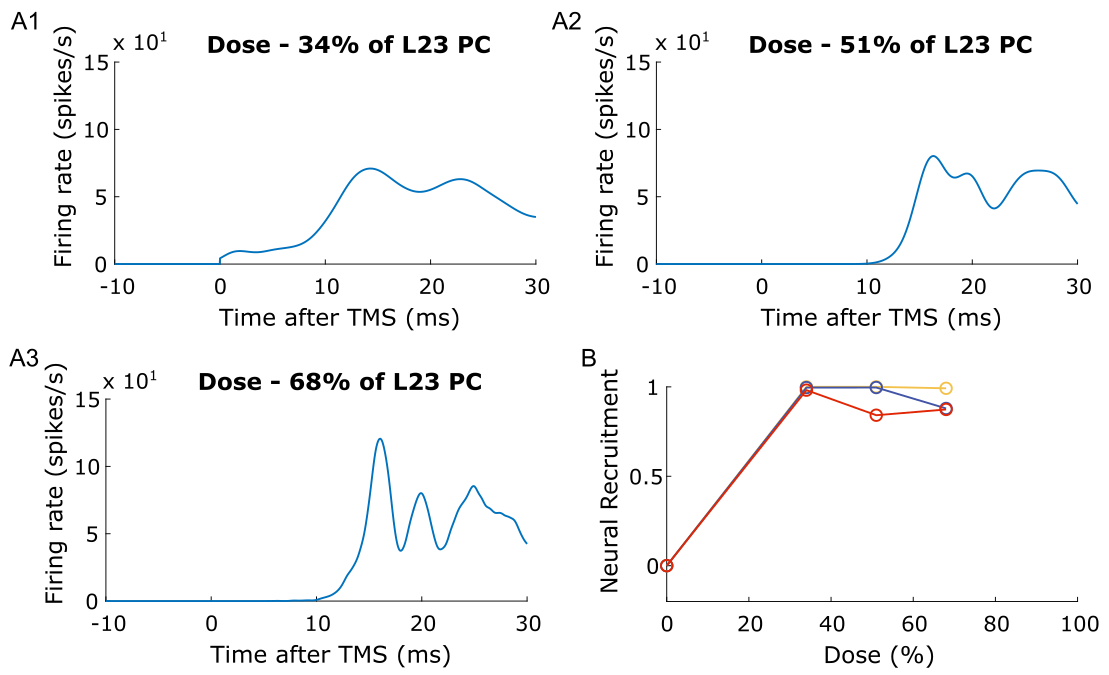

**Figure 5-1:** Response of L5 PNs, obtained by convolving their firing rates with a Gaussian function, to activation of different proportions of PNs in the S1-MC model. Although the S1-MC model generated rhythmic spiking activity, the response timing in the S1-MC model did not match the I-wave timing in experimental studies. Number of L5 PNs exhibiting rhythmic spiking activity peaks as a function of stimulation intensity in the S1-MC model. The maximum inter-peak interval to detect I-waves was 5 ms.

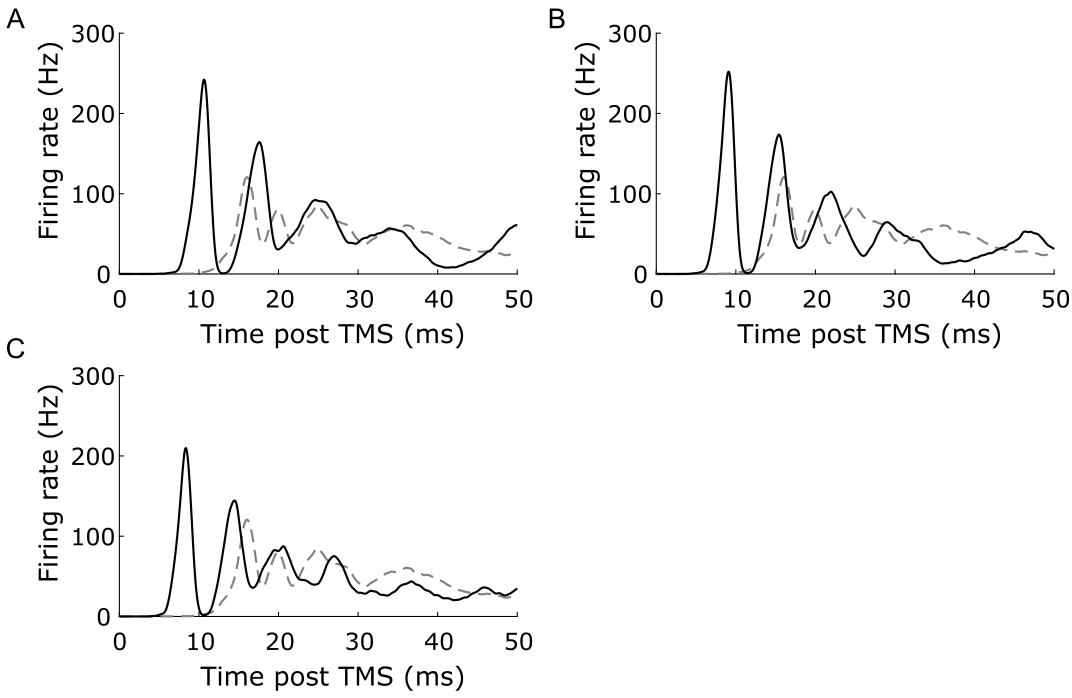

**Figure 6-1:** Response of L5 PNs, obtained by convolving their firing rates with a Gaussian function, at different levels (A) x2, (B) x3, (C) x4 of baseline AMPA synaptic conductance in the S1-MC model. The solid black trace indicates the response to AMPA synaptic modulation, while the control response is shown in the dashed black trace.

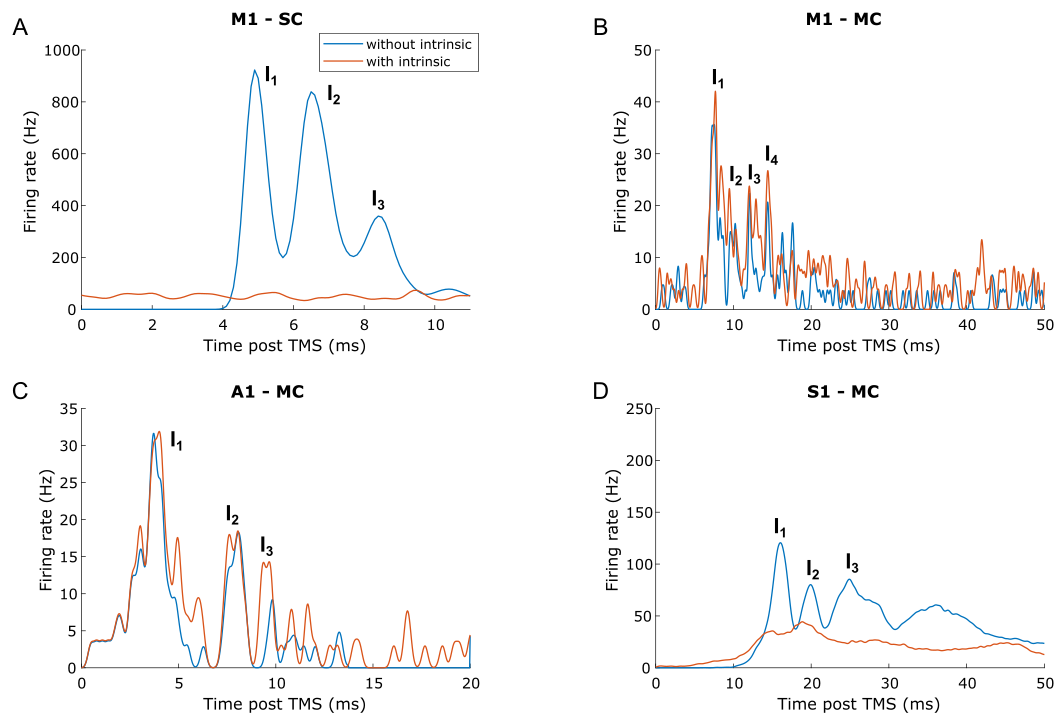

**Figure 6-2:** Response of L5 PNs, obtained by convolving their firing rates with a Gaussian function, with and without intrinsic activity for (A) M1-SC model, (B) M1-MC model, (C) A1-MC model and (D) S1-MC model.

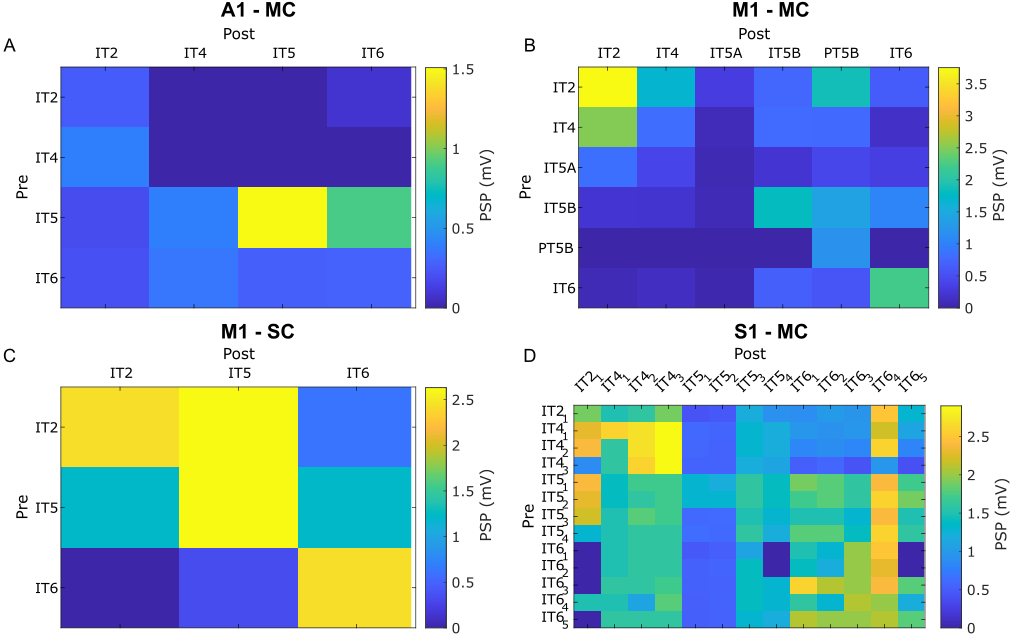

**Figure 9-1:** Post-synaptic potential (PSP) of monosynaptic excitatory connections in (A) A1-MC model, (B) M1-MC model, (C) M1-SC model, (D) S1-MC model.

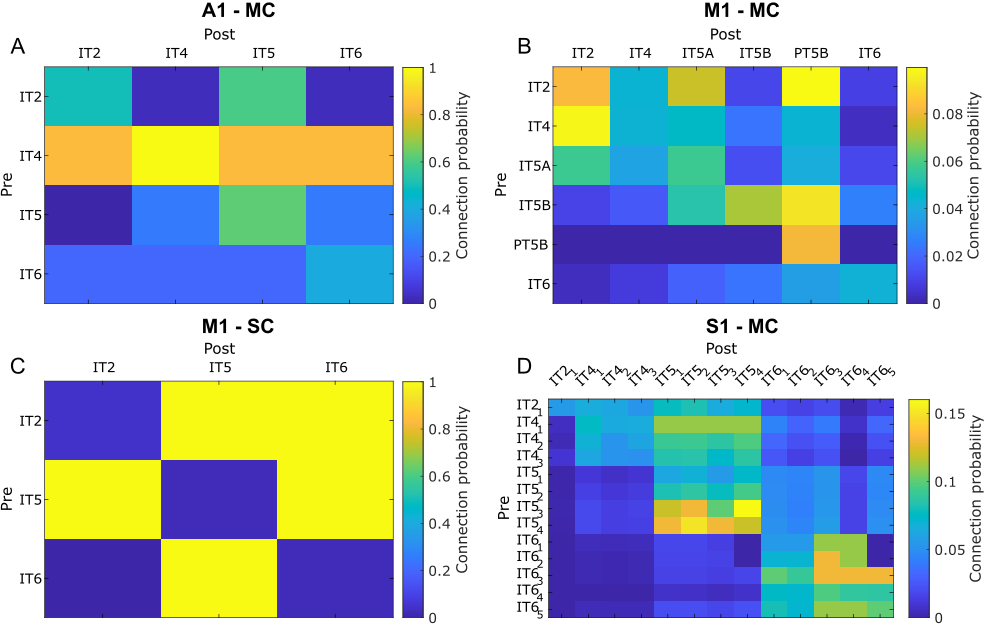

**Figure 9-2:** Connection probability of monosynaptic excitatory connections in (A) A1-MC model, (B) M1-MC model, (C) M1-SC model, (D) S1-MC model.
